# Selection for reducing energy cost of protein production drives the GC content and amino acid composition bias in gene transfer agents

**DOI:** 10.1101/2020.05.06.081315

**Authors:** Roman Kogay, Yuri I. Wolf, Eugene V. Koonin, Olga Zhaxybayeva

## Abstract

Gene transfer agents (GTAs) are virus-like elements integrated into bacterial genomes, particularly, those of Alphaproteobacteria. The GTAs can be induced under nutritional stress, incorporate random fragments of bacterial DNA into mini-phage particles, lyse the host cells and infect neighboring bacteria, thus enhancing horizontal gene transfer. We show that the GTA genes evolve under pronounced positive selection for the reduction of the energy cost of protein production as shown by comparison of the amino acid compositions with both homologous viral genes and host genes. The energy saving in GTA genes is comparable to or even more pronounced than that in the genes encoding the most abundant, essential bacterial proteins. In cases when viruses acquire genes from GTAs, the bias in amino acid composition disappears in the course of evolution, showing that reduction of the energy cost of protein is an important factor of evolution of GTAs but not bacterial viruses. These findings strongly suggest that GTAs are bacterial adaptations rather than selfish, virus-like elements. Because GTA production kills the host cell and does not propagate the GTA genome, it appears likely that the GTAs are retained in the course of evolution via kin or group selection. Therefore, we hypothesize that GTA facilitate the survival of bacterial populations under energy-limiting conditions through the spread of metabolic and transport capabilities via horizontal gene transfer and increase of nutrient availability resulting from the altruistic suicide of GTA-producing cells.

**Importance:** Kin and group selection remain controversial topics in evolutionary biology. We argue that these types of selection are likely to operate in bacterial populations by showing that bacterial Gene Transfer Agents (GTAs), but not related viruses, evolve under positive selection for the reduction of the energy cost of a GTA particle production. We hypothesize that GTAs are dedicated devices for the survival of bacteria under the conditions of nutrient limitation. The benefits conferred by GTAs under nutritional stress appear to include horizontal dissemination of genes that could provide bacteria with enhanced capabilities for nutrient utilization and the increase of nutrient availability through the lysis of GTA-producing bacteria.

## Introduction

Gene transfer agents (GTAs) are phage-like entities that are known to be produced by several groups of bacteria and archaea (1, 2). Unlike phages, GTAs do not package genes encoding their own structural proteins, and instead package pieces of DNA of the cell that produces them. The biological functions of the GTAs are not well understood, but the leading hypothesis is that GTAs are dedicated vehicles for horizontal gene transfer (HGT) (3, 4). The GTAs can be induced by stress (5) and, after packaging host DNA and lysing the host cell, can infect neighboring cells (1, 6). These cells can integrate the DNA contained within the GTAs, and thus can acquire new alleles, some of which could increase their fitness (7). GTAs are thought to have evolved from different viral ancestors on at least five independent occasions (2), and in Alphaproteobacteria, GTAs appear to have been maintained for many millions of years (8). Such convergent acquisition and long-term persistence of these elements suggests that GTAs provide a selective advantage for their host populations (2).

The best-studied GTA (RcGTA) comes from the alphaproteobacterium *Rhodobacter capsulatus* (9). Its production is directed by at least five loci that are scattered across the *R. capsulatus* genome, with 17 genes that encode most of the proteins necessary for the production of the RcGTA particles located in one locus (**Table S1**) (10). This locus, also known as the ‘head-tail’ cluster (2), is detectable in many alphaproteobacterial genomes (8, 11). Across Alphaproteobacteria, the RcGTA-like ‘head-tail’ clusters appear to evolve relatively slowly (1), have an elevated GC-content relative to the host genome (8), and have skewed amino acid composition when compared to their viral homologs (11).

Because bacteria and archaea occupy diverse ecological niches, they face different levels and directions of selective pressures and have different mutation rates, skewed GC-content and amino acid composition that emerged from multiple, intertwined processes. As a result, the genomic GC-content of bacterial and archaeal species varies in the wide range from less than 20% to more than 75% (12) and cannot be explained solely by the universal mutational AT-bias (13). Several studies have shown that the availability of different nutrients in the environment can act as a selective force and is involved in shaping the GC content of genomes and amino acid content of the encoded proteins. For example, inhabitants of nitrogen-poor environments tend to have a low content of G and C nucleotides and of amino acids containing nitrogen in their side chains (14, 15). Because A and T each contain one nitrogen atom less than G and C, respectively, the reduced usage of the G and C allows an organism to minimize the demand for the limiting nitrogen during replication and transcription. By contrast, carbon limitation could drive long-term elevation of the genomic GC-content (16, 17), likely, because small (carbon-poor) amino acids are preferentially encoded by GC-rich codons (18).

In addition to the GC-content fluctuation between species, there is also a considerable GC-content heterogeneity within single bacterial and archaeal genomes. For example, bacterial genomes can be subject to GC-biased gene conversion and thus recombination hotspots within a genome can have elevated GC-content compared to the rest of the genome (19). Also, highly expressed genes tend to have an elevated GC-content and, accordingly, their highly abundant protein products have a skewed amino acid composition (20). Because highly abundant proteins appear to be optimized for low cost of production (21, 22), the elevated GC-content of highly expressed genes can be explained by selection for GC-rich codons that tend to encode small, energetically cheap amino acids. Generally, molecular composition of genes and proteins appears to reflect various selection pressures, among which those associated with energy savings are prominent.

Thus, there are two possible explanations for the observed skew in both the GC-content and amino acid composition of the RcGTA-like genes and proteins. Under one scenario, selection and mutational biases act on the base composition, so that the amino acid bias is a byproduct of the skewed GC-content. Under the second scenario, selection could favor the skewed amino acid composition, resulting in a biased GC-content due to the structure of the genetic code. Here, we present evidence for the second scenario and show that the observed amino acid bias is driven by selection to reduce carbon utilization and biosynthetic cost of production of the RcGTA-like proteins. We show that the energy expense of the production of RcGTA-like proteins is comparable to that of the highly expressed housekeeping genes. For some of the amino acid changes, we identify clear signatures of positive selection towards amino acids with a smaller number of carbons in their side chains. We hypothesize that evolution of RcGTA-like elements was affected by selection to minimize cellular energy investment into their production under nutrient-poor conditions.

## Results

### Elevated GC-content in RcGTA-like regions is due to the higher GC-content in the first and second codon positions of the coding genes

Because of the degeneracy of the genetic code, GC3-content is known to track the overall GC-content of genomic regions (23). Hence, if the GC-content of RcGTA-like ‘head-tail’ clusters is elevated because they reside in GC-rich genomic regions, the GC-content in the third, primarily synonymous codon positions (GC3-content) of the RcGTA-like genes is expected to be higher compared to the genomic average of the GC3-content. Moreover, the elevated GC3-content would not be limited to the genes in the RcGTA-like region but would be apparent in the adjacent genes as well. To test this hypothesis, we examined homologs of one RcGTA locus (‘head-tail’ cluster) in 212 alphaproteobacterial genomes (see **Materials and Methods**) (8, 11). Although we analyzed homologs of only one locus from one GTA only, for brevity, we hereafter refer to these regions simply as “GTA regions”, and to genes and encoded proteins in these regions as “GTA genes” and “GTA proteins”. Contrary to the aforementioned expectation, we found no significant difference between the GC3-content of GTA genes of the 212 alphaproteobacterial genomes, their neighboring genes and all genes in the genome (Kruskal-Wallis H test, p-value = 0.62; **Figure 1**). By contrast, the GC1- and GC2-content of GTA genes are significantly higher than the corresponding values for both the neighboring genes (Dunn’s test, p-value < 0.0001) and the genes across the entire genome (Dunn’s test, p-value < 0.0001) (**Figure 1**). Furthermore, the genes adjacent to the GTA regions do not have elevated GC1- and GC2-content when compared to the genes in the entire genome (Dunn’s test, p-value = 1), indicating that the elevated GC1- and GC2-content is limited to the GTA genes. Due to the relationship between codons and amino acids in the genetic code, the elevated GC1- and GC2-content of an open reading frame (ORF) translates into a biased amino acid composition of the encoded protein. Indeed, a significant amino acid composition bias in the GTA proteins has been demonstrated previously (11). Specifically, the relative abundance of amino acids encoded by GC-rich codons is significantly higher in the GTA genes than the genomic average (**Figure S1;** Student’s t-test, p-value < 0.0001; See **Materials and Methods** for definition of GC-rich codons). Taken together, these findings suggest that the GC-content of GTA regions in Alphaproteobacteria is driven by selection for a specific amino acid composition of the encoded proteins.

**Figure 1.**
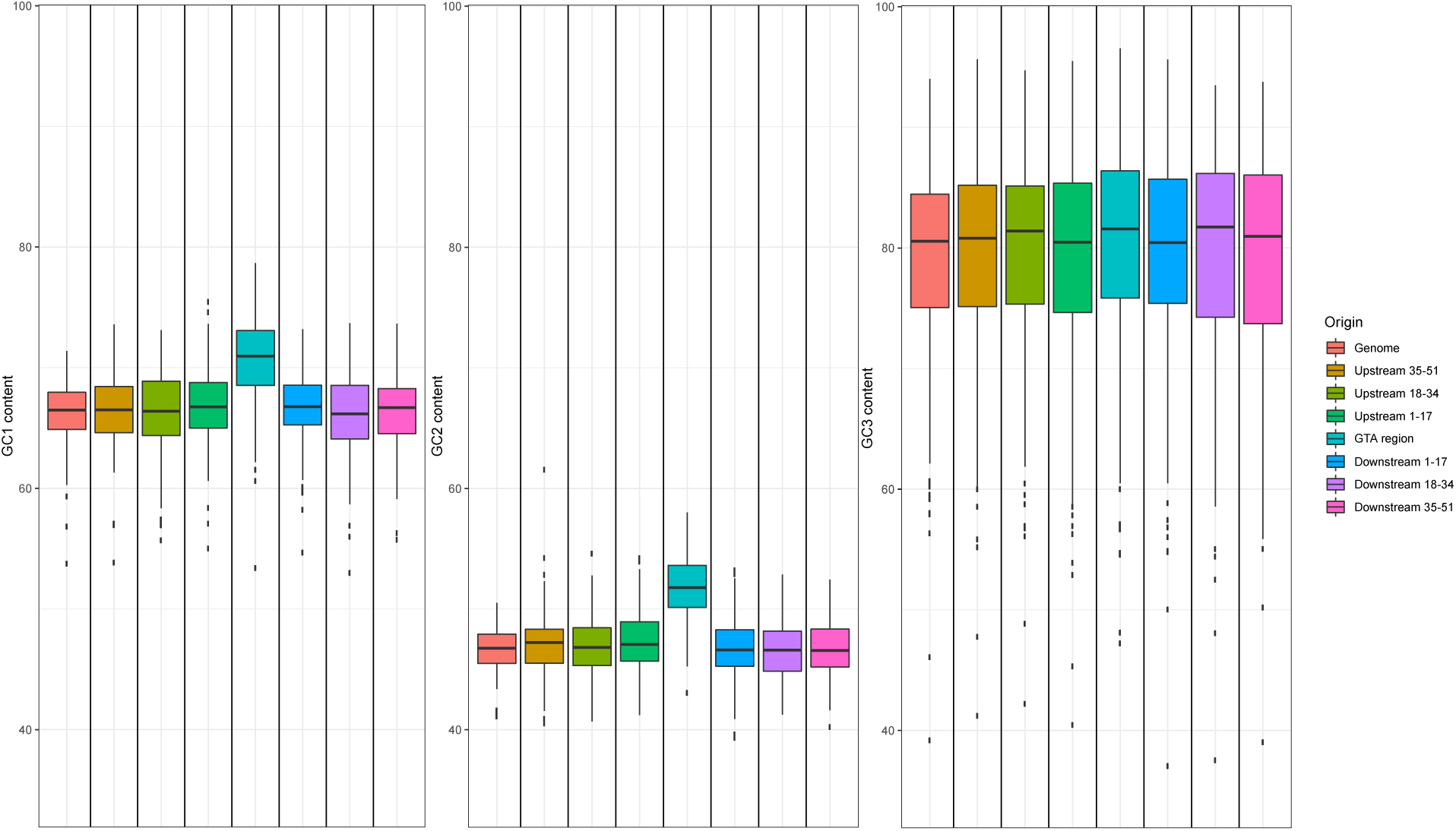
The GC1-, GC2- and GC3-content of GTA regions, their immediate neighborhoods and all protein-coding genes in 212 alphaproteobacterial genomes. The neighborhoods immediately upstream and downstream of a GTA region consists of 17 genes each. Boxplots represent median values bounded by the first and third quartiles. Whiskers show the values that lie in the range of 1.5*interquartile rule. Dots outside of the whiskers are the outliers.

### Proteins encoded in GTA regions contain smaller number of carbons and are energetically cheaper than their viral homologs

The RcGTA production has been experimentally demonstrated to be stimulated by carbon depletion (5). Furthermore, knockout of the RcGTA-like genes in three alphaproteobacterial strains (24) resulted in a significant decrease in fitness of the mutants under growth conditions with alternative carbon sources that might not be utilized by these strains (11). If GTAs are indeed produced under conditions of limited carbon availability, the observed amino acid bias in the GTA genes might represent an adaptation in the GTA-containing lineages to utilize energetically cheaper amino acids for GTA particle production. To test this hypothesis, we compared the number of carbons in amino acid side chains and costs of amino acid biosynthesis (measured as the number of high-energy phosphate bonds) in GTA proteins and by their viral homologs. We assumed that (a) all amino acids are produced by bacteria *de novo*, as at least 174 of the analyzed genomes can produce 19 or all 20 amino acids (**Figure S2**), and (b) viral infections are not specifically associated with the carbon-limited conditions, and therefore, viral homologs of RcGTA genes should not be subject to selection for energy saving. Consistent with the proposed hypothesis, for all of the 12 genes with sufficient number of viral homologs to estimate statistical significance (**Table S1**), GTA proteins have both a significantly smaller number of carbons (Mann-Whitney U test, all 12 Bonferroni-corrected p-values < 0.01; **Figure 2A**) and a significantly reduced cost of amino acid biosynthesis than their viral homologs (Mann-Whitney U test, all 12 Bonferroni-corrected p-values < 0.01; **Figure 2B**).

**Figure 2.**
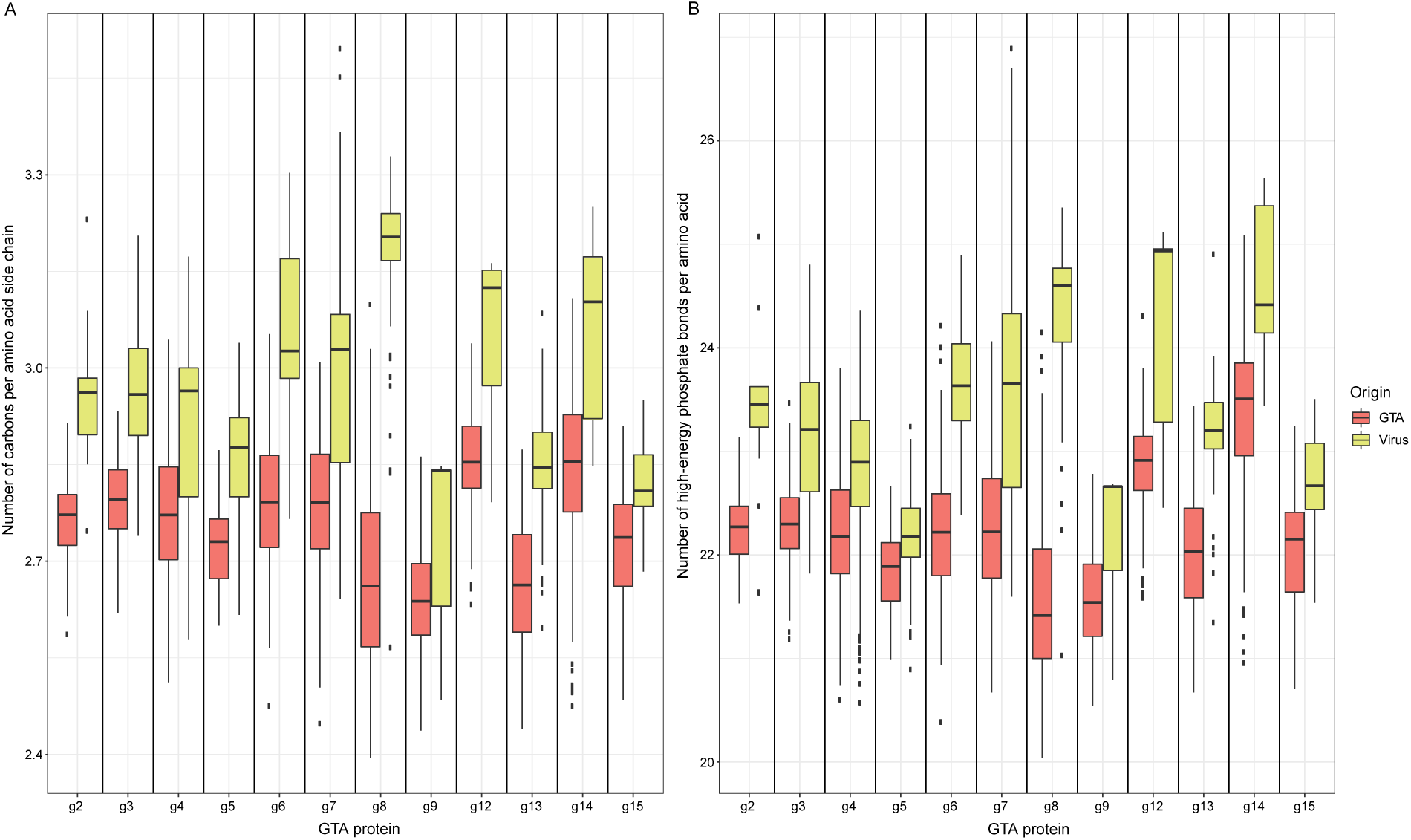
Carbon content (A) and biosynthetic cost (B) of proteins encoded by GTA genes in 212 alphaproteobacterial genomes and their viral homologs. Boxplots represent median values that are bounded by the first and third quartiles. Whiskers show the values that lie in the range of 1.5*interquartile rule. Dots outside of the whiskers are the outliers. The number of data points in each boxplot is listed in **Table S1**.

To demonstrate that the observed differences in the carbon content of the GTA and viral proteins are not simply due to the compositional bias present in the ancestor of the alphaproteobacterial GTA elements (8), we sought to examine only a subset of viral homologs that are presumed to be horizontally acquired from the GTA regions. Genes with significant sequence similarity to GTA genes have been previously found in viruses and inferred to be horizontally acquired from GTAs on the basis of phylogenetic reconstruction (10, 25). In our phylogenetic analyses, we examined several viral genes of this apparent origin (**Table 1, Figure S3**; also see **Materials and Methods** for details). Under the assumption of no selection for energy saving in viruses, we expect the carbon content of the GTA genes acquired by viruses to increase after their relocation to the virus genomes. Indeed, in all cases, the carbon content of the now-viral homologs consistently (and, overall, significantly) increased compared to the inferred ancestral state at the time of acquisition (**Table 1, SI Figure S3**).

**Table 1.** Change in the carbon content between viral homologs of the GTA proteins and their closest GTA ancestral node.

| **GTA gene** | **Virus name** | **Change in the number of carbons per side chain of an amino acid** | **p-value** | **Alignment length** |
| --- | --- | --- | --- | --- |
| *g6* | *Cellulophaga* phage phi10 1 | +0.605 | <0.001 | 193 |
| *g7* | *Cellulophaga* phage phi18 1 | +0.394 | 0.001 | 147 |
| *g7* | *Streptomyces* phage phiSASD1 | +0.167 | 0.179 | 147 |
| *g7* | *Salmonella* phage ST64B | +0.222 | 0.048 | 147 |
| *g7* | *Salmonella* phage 118970 sal3 | +0.229 | 0.042 | 147 |
| *g7* | *Shigella* phage SfIV | +0.184 | 0.115 | 147 |
| *g7* | Enterobacteria phage SfV | +0.244 | 0.083 | 147 |
| *g7* | *Shigella* phage SfII | +0.191 | 0.107 | 147 |
| *g10* | *Rhizobium* phage 16-3 | +0.105 | 0.271 | 123 |
| *g12* | *Rhodobacter* phage RcCronus | +0.123 | 0.081 | 228 |
| *g13* | *Paracoccus* phage vB PmaS R3 | +0.048 | 0.226 | 304 |
| *g13* | *Dinoroseobacter* phage vB DshS R5C | +0.027 | 0.383 | 304 |
| *g13* | *Roseobacter* phage RDJL Phi 1 | +0.005 | 0.447 | 304 |
| *g13* | *Roseobacter* phage RDJL Phi 2 | +0.019 | 0.388 | 304 |
| *g14* | *Rhodobacter* phage RcRhea | +0.191 | 0.108 | 166 |
| *g15* | *Rhodobacter* phage RcRhea | +0.147 | <0.001 | 1369 |
| *g15* | *Rhodobacter* phage RcCronus | +0.143 | <0.001 | 1369 |
| *Cumulative across 7 genes* |  | +0.163 | <0.001 | 2530 |

### Energetic cost of the GTA proteins is as low as that of essential bacterial proteins

Highly expressed genes have been demonstrated to evolve under selection to decrease the energetic cost of the encoded protein production (20). Indeed, 20 single-copy housekeeping genes involved in translation ([J] COG category; (26)) (**Table S2**), and therefore presumed to be expressed at relatively high levels under any conditions, collectively, have a significantly lower energetic cost than the average of all proteins encoded in a genome, as measured by both side chain carbon utilization and biosynthetic cost of production per amino acid (**Figure S4;** Mann-Whitney U test, p-values < 0.0001). The biosynthetic cost per amino acid of the GTA proteins was found to be statistically indistinguishable from that of the products of the 20 highly expressed genes (α of 0.01; Mann-Whitney U test, p-value = 0.3372), and remarkably, utilize even less carbon (α of 0.01; Mann-Whitney U test, p-value < 0.0001) (**Figure S4**).

### Reduction in carbon utilization varies among GTA genes and across bacterial taxa

To investigate how reduction of carbon content evolved from the common ancestor of the examined GTA genes to the extant forms, we reconstructed the number of carbons per amino acid at the ancestral nodes of individual evolutionary trees of 14 GTA genes (those with at least one detectable viral homolog; **Table S1**). To correct for differences in the GC-content across taxa (which affects the carbon content of the encoded proteins), for each taxon we normalized the number of carbons per amino acid of GTA proteins by that of 26 housekeeping proteins (**Table S2**). No unifying pattern of directional selection towards the lower carbon content was detected across all genes and all taxa (**Figure S5**). This lack of an overall signal was not surprising because GTA genes can be horizontally transferred across taxa (8), have different evolutionary rates among and within taxa (8), and are likely to reach unequal translation levels during GTA production (27). These differences would make the carbon content optimization gene- and taxon-specific, blurring the net effect. However, members of the order *Sphingomonadales* show the most pronounced reduction in carbon utilization for the GTA regions overall, as well as for the majority of individual genes (**Figure 3**). Notably, many *Sphingomonadales* species can live under nutrient-depleted conditions (28).

**Figure 3.**
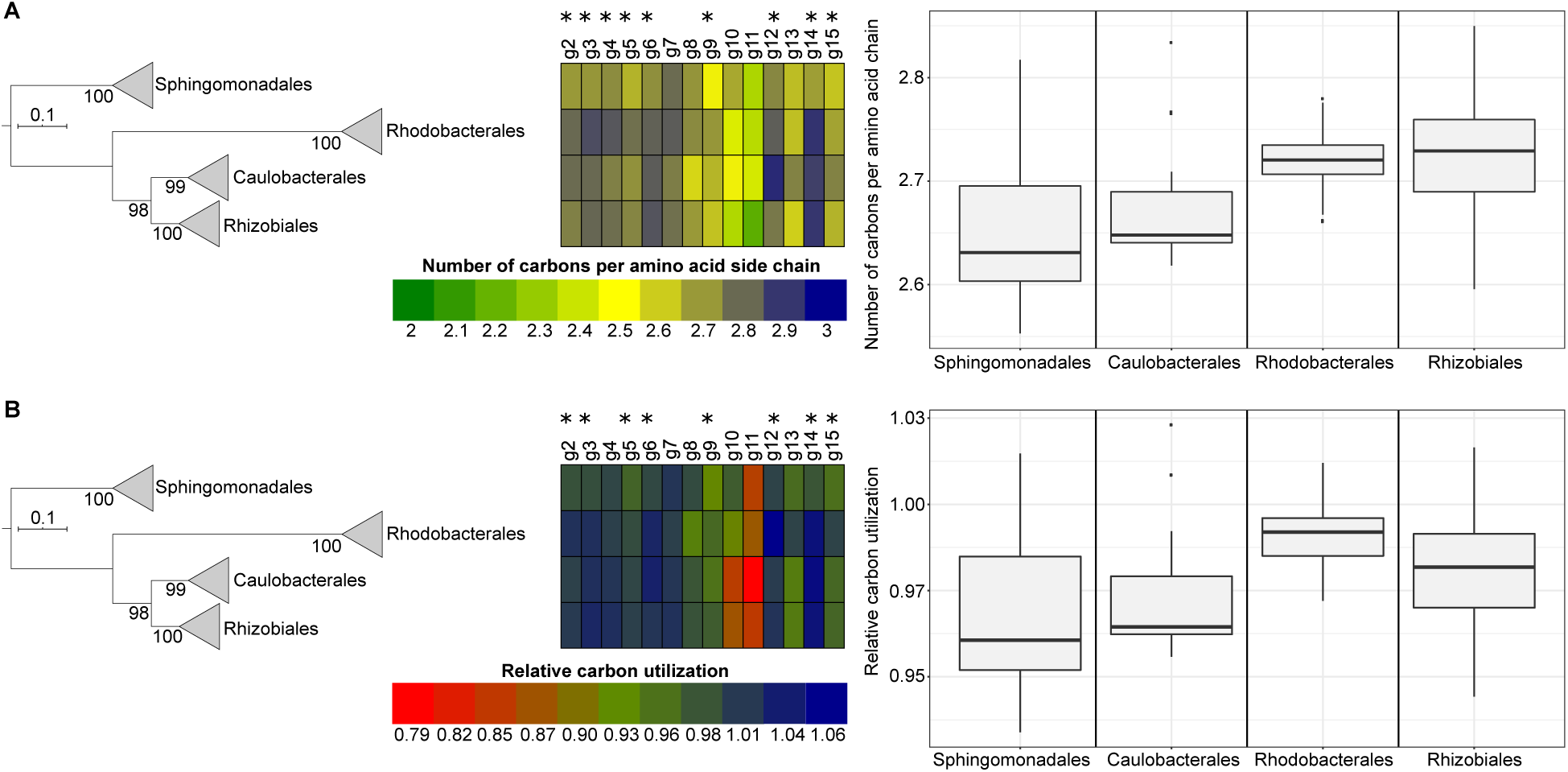
Carbon content of GTA proteins for four orders of the class *Alphaproteobacteria*. For each GTA protein, the heatmap visualizes the number of carbons per the side chain in amino acid averaged across taxonomic order. The numbers are shown either as raw values (**panel A**), or as values normalized by the carbon content of proteins encoded by 26 single-copy genes (**panel B**). The asterisks mark GTA proteins with significantly lower numbers of carbons per amino acid in the *Sphingomonadales* order than in the other three orders combined (α of 0.01; Mann-Whitney U test, all p-values < 0.01). Boxplots summarize the distribution of carbon content within each alphaproteobacterial order averaged across the examined GTA genes. Median values are bounded by the first and third quartiles. Whiskers show the values that lie in the range of 1.5*interquartile rule and dots outside of the whiskers are the outliers. The phylogenetic tree is the reference alphaproteobacterial phylogeny (see **Materials and Methods** for details), in which branches are collapsed at the taxonomic rank of order. Numbers at the tree nodes represent bootstrap support values. Scale bar, number of substitutions per site.

### In *Sphingomonadales*, the decrease in carbon content of GTA proteins is driven by positive selection

To evaluate whether diversifying (positive) selection plays a role in the observed reduction of carbon utilization in the GTA genes in *Sphingomonadales*, we tested for evidence of positive selection in individual sites on the branch leading to this clade. For 9 of the 14 evaluated genes, the model of positive selection on the branch was a significantly better fit than the neutral null model (**Table S3**). For 8 of these 9 genes, members of the *Spingomonadales* clade showed significant decrease in the carbon utilization relative to three other orders (Mann-Whitney U Test; α of 0.01, p -values < 0.01; **Table S4**; **Figure 3**). Conversely, for 4 of the 5 genes that did not show evidence of positive selection, there was no significant decrease in the carbon content of proteins in the *Sphingomonadales* genomes (**Figure 3**).

To assess how the specific sites that are inferred to be subject to positive selection contribute to the carbon content of the *Sphingomonadales*’ GTA genes, we examined carbon content of amino acids in the sites with >0.95 posterior probability of being subject to positive selection. For 8 of the 9 positively selected genes, these sites substantially contributed to the decrease in carbon utilization in *Sphingomonadales* (**Table 2, Table S5**). This trend is manifested, in particular, by the observed replacements of aromatic amino acids, which contain relatively high numbers of carbons and have excessive biosynthetic costs, with non-aromatic amino acids (**Figure S6**). The observed replacements of tryptophan with phenylalanine indicate that, under a constraint of maintaining an amino acid with similar physicochemical properties, there is selection for utilization of a cheaper amino acid (**Figure S6**). Mapping of the positively selected sites in the *Sphingomonadales*’ g5 homolog onto a structural model of the T5 bacteriophage major capsid protein shows that these sites tend to be located on the surface of the protein (**Movie S1**). This example suggests that carbon-saving replacements preferentially occur in sites that are not involved in the folding of GTA proteins, allowing the GTAs to preserve the functionality of their proteins at reduced production costs.

**Table 2.** Contribution of positively selected sites to the reduction of carbon utilization in GTA proteins of *Sphingomonadales*.

| GTA protein | Number of sites under the positive selection | Average change in number of carbons by the contribution of all sites under the positive selection | Number of sites that contribute to the decrease in number of carbons |
| --- | --- | --- | --- |
| g2 | 13 | -0.22 | 6 |
| g3 | 33 | -0.72 | 22 |
| g4 | 29 | -0.42 | 13 |
| g5 | 12 | -0.39 | 8 |
| g6 | 11 | +0.16 | 5 |
| g9 | 29 | -0.68 | 16 |
| g12 | 23 | -0.52 | 13 |
| g13 | 31 | -0.44 | 16 |
| g15 | 27 | -0.55 | 15 |

## Discussion

We show here that the elevated GC-content of GTA regions is driven by selection towards encoding proteins with energetically cheaper amino acids. Although GC-rich genes have an increased cost of mRNA expression, cells spend much more energy on the synthesis of amino acids than on the synthesis of ribonucleotides (20, 29). Hence, the elevation of GC-content in non-synonymous codon positions (GC1 and GC2) reduces the energetic expenses on the production of the respective proteins. Consistent with this notion, energy savings for GTA proteins are as pronounced or even greater than those for highly expressed housekeeping genes that are known to utilize cheaper and smaller amino acids (20). Given that production of RcGTA-like particles in Alphaproteobacteria occurs in the stationary phase (2, 30) and is associated with carbon depletion (5, 11), the shift in GC-content of GTA genes and amino acid composition of their products likely reflects the adaptation for their efficient expression under such conditions.

The change in the amino acid composition of GTA proteins was not uniform across the examined alphaproteobacterial lineages. These differences are not unexpected because GTA-carrying bacteria live in different environments and under different selection pressures. We demonstrated that, on the branch leading to *Sphingomonadales*, the decrease in carbon content of the GTA proteins is driven by positive selection for the use of cheaper amino acids. We hypothesize that the last common ancestor of *Sphingomonadales* evolved in a nutrient-depleted environment that selected for the reduction in the use of energetically expensive amino acids in the GTA proteins.

Although bacterial viruses also spend disproportionate amounts energy on translation (31), our analysis of viral genes that apparently were acquired by viruses from bacterial GTAs shows a decrease in GC1 and GC2 content, with the concomitant increase in protein production energy cost. Thus, positive selection for cost saving, probably ceases to substantially affect the evolution of these genes once they are transferred to virus genomes. Lytic bacteriophages reproduce rapidly, with a typical burst size of about 200 virions that hijacks about 30% of the host energy budget (31). Under the conditions of such brief, explosive growth, energy saving might not be an important selective factor. Differences in the viral burst sizes imply that selection for energy saving could play some role. However, such selection is expected to be weak due to other constraints affecting the lytic viruses, such as fluctuations in the host energy budget, often error-prone viral replication machinery, and the main evolutionary pressure being evasion of host defense systems (32, 33). Thus, our observations provide additional evidence that GTAs are not selfish, virus-like agents but rather microbial adaptations.

Taken together, our findings, and in particular, the evidence of positive selection for energy saving in *Sphingomonadales*, are in line with the previous suggestions that maintenance of GTAs and production of GTA particles confers some advantage to the bacterial hosts (2, 7). Because GTA-producing cell lyses, the reduction of energy utilization for the production of GTA particles has to be beneficial at the population or community level, that is, it needs to involve some form of kin or group selection (34, 35). The nature of such benefit(s) is not entirely clear, but it appears likely that the GTAs, effectively, are devices for survival under energy- or nutrient-limited conditions that are common in bacterial ecology. More specifically, GTAs could provide two types of adaptations. Previous studies suggest that oligotrophic conditions do not interfere with the capacity of bacteria to engage in genetic exchange (36). Moreover, the nutrient limitation can upregulate horizontal gene transfer via transformation (37), suggesting potential benefits of gene exchange under adverse conditions of energy or nutrient limitations. Conceivably, HGT mediated by the GTAs can confer additional metabolic or transport capacities to the recipient bacteria. Additionally, GTAs could be perceived as a mechanism bacterial programmed cell death (38, 39). Under this type of adaptation, the GTA-mediated lysis of a fraction of the bacterial community would decrease the population density and increase the nutrient availability per cell, by supplying additional nutrients released from the lysed cells.

## Materials and Methods

### Generation of GTA and viral datasets

The initial dataset of 422 GTA regions in 419 alphaproteobacterial genomes consisted of 88 regions identified by Shakya et al (8) and 334 regions in complete alphaproteobacterial genomes predicted by Kogay et al. (11). Four GTA regions from the *Methylobacterium nodulans* ORS2060 genome were removed due to their questionable assignment as GTAs (8). Because our previous GTA prediction procedure (11) screened for the presence of only 11 of the 17 homologs of the RcGTA head-tail cluster (1), the remaining 6 homologs were identified using BLASTP (40) (version 2.6.0, e-value = 0.1, manually curated homologs from Kogay et al. (11) as queries), with subsequent restriction of the hits to the regions with previously identified GTA genes. To reduce the computational cost of the downstream analyses, highly similar GTA regions were excluded. To this end, genomes that contained the 418 GTA regions were clustered into OTUs using furthest neighbor clustering and Average Nucleotide Identity (ANI) cutoff of 95%. The ANI values were calculated using fastANI v.1.1 (41). From each of the identified 215 OTUs, only the GTA region with the largest number of the relevant genes was retained. Further removal of the regions that contained less than 9 genes resulted in the final dataset of 212 GTA regions.

To obtain viral homologs of the GTA genes, genes from the 212 GTA regions were used as queries in BLASTP searches (40) (version 2.6.0, e-value = 0.001, query and subject coverage of at least 60%) against the viral RefSeq database (release 96, accessed on October 2019) (42).

The numbers of identified alphaproteobacterial and viral homologs for the 17 RcGTA genes are shown in **Table S1**.

### Calculation of GC-content for the 212 alphaproteobacterial genomes

The GTA region’s neighborhood was defined as 51 genes upstream and 51 genes downstream of the region. Each neighborhood was divided into 6 non-overlapping regions with 17 genes each. For each neighborhood region, the GTA region, and all annotated genes in the genome, GC1-, GC2-, and GC3-content values were calculated using an in-house script. The significance of the GC-content differences among the obtained 8 groups was assessed using the Kruskal-Wallis H test followed by the Dunn’s test (43). The p-values were adjusted for multiple testing using the Bonferroni correction.

### Calculation of the relative abundance of amino acids encoded by GC-rich codons for 212 alphaproteobacterial genomes

The amino acids that are encoded by GC-rich codons were defined as those that have G or C in the first and second codon positions (alanine, arginine, glycine and proline). For each genome, the amino acid frequencies were calculated for the pooled set of proteins encoded by genes in the GTA region, as well as for the pooled set of proteins encoded by all genes in a genome. The significance of the difference in relative abundances of the 4 amino acids encoded by GC-rich codons in the two sets was assessed using the Student’s t-test.

### Calculation of carbon content and biosynthetic cost of amino acids in the encoded proteins

Because differences in the carbon-content of amino acids are determined solely by the composition of their side chains, for each amino acid sequence encoded by a GTA gene (or its viral homolog), the number of carbons in the side chains of the amino acids was counted and normalized by the length of the encoded polypeptide. Additionally, for each amino acid sequence encoded by a GTA gene (or its viral homolog), the average biosynthetic cost of protein production per amino acid, defined as the number of high-energy phosphate bonds needed to produce a particular amino acid, was calculated. Because almost all of the 212 alphaproteobacteria containing the GTA regions are either obligate or facultative aerobes, the individual costs of amino acid production already computed for *Escherichia coli* by Akashi and Gojobori (44) were used. The significance of the differences in the carbon utilization and biosynthetic cost between GTA proteins and viral homologs was assessed using the Mann-Whitney U test, followed by the Bonferroni correction of p-values to account for multiple testing.

### Verification of amino acid biosynthesis pathways in the alphaproteobacterial genomes

Presence of the amino acid biosynthesis pathways in the genomes was evaluated using the KEGG database release 92 (45). For 189 of the 212 alphaproteobacteria, either its own genome (186 genomes) or the genome of a close relative (ANI > 95%; 3 genomes) were examined. For the remaining 23 genomes, no information from the closely related genomes was available in KEGG. For each of the 189 genomes, the map of amino acid biosynthesis (map number = 01230) was examined for completeness. If key enzymes were missing, additional maps (map number = 00250 – 00400) were evaluated to identify alternative enzymes that could catalyze the same reactions. If alternative enzymes were not found, *Escherichia coli* homologs that catalyze the missing steps were used as queries for a BLASTP search of the genome (version 2.6.0, e-value 0.001, query coverage of at least 50%) and the RefSeq annotations of the obtained matches were examined. If a complete biosynthetic pathway of an amino acid could not be reconstructed, the genome was designated as “auxotrophic” for the biosynthesis of the given amino acid.

### Exclusion of divergent viral homologs

To minimize possible misplacement of viral homologs due to long branch attraction, we have identified and excluded divergent viral homologs using the following procedure. Amino acid sequences of GTA genes and their viral homologs were aligned using MAFFT v 7.305 with the ‘auto’ setting (46). Phylogenetic trees from individual gene alignments were reconstructed in the IQ-TREE v 1.6.7 (47) using the best substitution model detected by ModelFinder (48). The obtained trees were used as guides for the reconstruction of more accurate trees, using the profile mixture model “LG+C60+F+G” and the site-specific frequency models that were approximated by the posterior mean site frequency (49), as implemented in IQ-TREE.

To exclude viral homologs that are not closely related to GTA genes, only viral homologs nested within the taxonomic rank of alphaproteobacterial order with ultrafast bootstrap support >= 60% (1,000 pseudoreplicates; (50)) were retained. Because, for genes *g3, g4* and *g8*, large numbers of viral homologs were retained, only top 5 non-identical viral proteins most closely related to the alphaproteobacterial homologs were kept. The retained viral homologs were realigned with the GTA genes, and the phylogenetic trees were reconstructed and examined as described above. The process was repeated until all retained viral homologs grouped within alphaproteobacterial orders.

### Reconstruction of ancestral amino acid sequences

Amino acid sequences of the ancestral nodes of the reconstructed phylogenetic trees were reconstructed using FastML v 3.11 (51). Indels in the ancestral sequences were inferred using the maximum likelihood and probability cutoff of 0.5. Ancestral amino acid states of non-gapped states were determined using marginal reconstruction under LG substitution matrix (52), with heterogeneity in substitution rates among sites modeled using Gamma distribution (53).

### Reconstruction of the alphaproteobacterial reference phylogeny

In each of the 212 genomes containing GTA regions, 31 phylogenetic markers were detected and retrieved using AMPHORA2 (54). Amino acid sequences of these markers were aligned using MAFFT v 7.305 with the ‘auto’ setting (46). The best substitution matrix for each gene was determined using the *ProteinModelSelection.pl* script obtained from https://github.com/stamatak/standard-RAxML/tree/master/usefulScripts (last accessed November 2019). The individual gene alignments were concatenated, and each gene was treated as a separate partition (55) in the subsequent phylogenetic reconstruction. The maximum likelihood tree was reconstructed by the IQ-TREE v 1.6.7 (47), and Gamma distribution with four categories was used to account for heterogeneity in substitution rates among sites (53). Although no outgroup sequences were included into the alignment, for presentation purposes, the tree was rooted to reflect the branching of Alphaproteobacteria as previously observed (11).

### Retrieval of selected single-copy and highly-expressed genes

Twenty-six of the 120 phylogenetically informative genes (56) were found to be present in a single copy in all 212 genomes (**SI Table S2**). The 26 genes were extracted from each genome using hmmersearch v 3.1b2 via modified scripts from AMPHORA2 (54). The functional annotations of the 26 genes were examined using the eggNOG-mapper (57) based on the eggNOG orthology database v. 4.5 (58). Twenty of the 26 genes belong to the [J] COG category (“Translation, ribosomal structure and biogenesis”) and therefore were designated “highly-expressed” genes.

### Calculation of carbon utilization in extant and ancestral GTA genes

The relative carbon utilization of each extant protein encoded by a GTA gene was defined as the ratio of the average number of carbon atoms per site to that of the 26 single-copy genes. To calculate carbon utilization for the ancestral states, amino acid sequences of 14 GTA proteins with at least one viral homolog were aligned by MAFFT v 7.305 with the “auto” setting (46), and phylogenetic trees were reconstructed using IQ-TREE v 1.6.7 (47) using the best substitution model detected with ModelFinder (48). Using reconstructed phylogenies and carbon utilization data for extant proteins, carbon utilization at the internal nodes was inferred using the marginal maximum likelihood reconstruction, as implemented in the *phytools* package (59). The change of carbon utilization along the tree branches was deduced via equation 2 of Felsenstein (60), also as implemented in the *phytools* package (59).

To assess the significance in the increase of carbon content of the selected viral proteins in comparison to their inferred ancestral protein, for each of the seven GTA genes with such viral homologs, amino acid sequences of these extant viruses and their closest inferred ancestral sequence were retrieved and aligned via MAFFT using “linsi” settings (46). For each gene alignment, 1000 bootstrap replicates were generated in RAxML v 8.2.11 (61). For each bootstrap replicate, the net change in the number of carbons per amino acid between the viral protein and the ancestral protein was calculated. The p-value was defined as the proportion of bootstrap replicates with a zero or negative net change in the number of carbons per amino acid. Additionally, the cumulative net change in the number of carbons per amino acid across all 7 GTA proteins (**Table 1**) was calculated by adding up the net changes across individual genes. For genes with more than one viral homolog, the viral homolog with the smallest difference in the number of carbons per amino acid was selected to obtain a conservative estimate. The p-values were calculated as they were for individual comparisons.

### Detection of positive selection on the branch leading to *Sphingomonadales*

Using the phylogenetic trees and amino acid sequence alignments of the GTA proteins (see “**Calculation of carbon utilization states in contemporary and ancestral GTA genes”** section), evidence of episodic events of positive selection in the *Sphingomonadales* clade was inferred under the branch site A model, as implemented in the *codeml* package of PAML version 4 (62). Codon alignments of nucleotide sequences were obtained using *pal2nal* (63). The branch lengths in the corresponding phylogenetic trees were re-estimated in PAML. Because *g12* and *g15* genes vary in length between *Sphingomonadales* and other alphaproteobacterial orders, codons that were present in less than 50% and 80% of sequences in *g12* and *g15* datasets, respectively, were removed. For the null model (no positive selection), ω2a and ω2b were fixed to 1, and the significance for the alternative model (positive selection) was tested using the likelihood ratio test with one degree of freedom and α of 0.01. P-values were adjusted for multiple testing using the Bonferroni correction. A site was classified as being “under positive selection” if it had the probability of at least 0.95 in the Bayes Empirical Bayes estimation (64), and was present in at least of 50% of the *Sphingomonadales* branches and 50% of the remaining branches.

### Visualization of positively selected sites on the 3D model of capsomer

The amino acid sequences of the RcGTA genes were used in a BLASTP search (e-value < 0.01, low-complexity masking, and query coverage of at least 50%) against the PDB database (65) (last accessed November 2019). Only the *g5* gene query returned significant matches to the PDB database. The amino acid sequence of the top-scoring match (PDB ID – 5TJT) was retrieved and aligned with the representative *g5* homolog from *Sphingomonadales* (*Sphingobium amiense DSM 16289*) using the Needleman-Wunsch algorithm (66). Of the 12 sites classified as being under positive selection in the *Sphingobium amiense DSM 16289* homolog, 2 sites did not have homologous positions in the 5TJT sequence. The remaining 10 sites were mapped onto the 5TJT PDB structure using PyMol version 2.3 (The PyMOL Molecular Graphics System, Version 2.0 Schrödinger, LLC.)

## Data Availability

List of accession numbers of 212 alphaproteobacterial genomes with GTA regions, amino acid sequences of identified GTA proteins in alphaproteobacteria and viruses, as well as sequence alignments and phylogenetic trees used in the described analyses have been deposited to FigShare under the doi: 10.6084/m9.figshare.12071223.

## Acknowledgments

This work was supported in part by the National Science Foundation [NSF-DEB 1551674 to O.Z.]; by the Simons Foundation Investigator in Mathematical Modeling of Living Systems [327936 to O.Z.]; and by Intramural Research Program of the National Institutes of Health of the USA (National Library of Medicine), to YIW and EVK.

## Supplementary materials for this manuscript include the following

**Figures S1 to S6**

**Tables S1 to S5**

**Movie S1**

**Data in the FigShare repository** at doi 10.6084/m9.figshare.12071223.

## Supplementary Figure Legends

**Figure S1. The relative abundance of amino acids encoded by GC-rich codons in all proteins in 212 alphaproteobacterial genomes and in proteins from GTA regions.** Boxplots represent median values that are bounded by the first and third quartiles. Whiskers show the values that lie in the range of 1.5*interquartile rule. Dots outside of the whiskers represent the outliers.

**Figure S2. The reference phylogeny of 212 analyzed alphaproteobacterial genomes and presence/absence of amino acid biosynthetic pathways in these genomes.** The phylogenetic tree represents the reference phylogeny of the analyzed genomes (see **Materials and Methods** for details). The presence (in green) and absence (in blue) of amino acid biosynthetic pathways in a genome is shown next to the taxon name of the genome. For 23 genomes with pathway data shown in black, no pathway information was available in the KEGG database. Scale bar, number of substitutions per site.

**Figure S3. Carbon content and GC content of proteins and genes, respectively, from GTAs and select viral homologs mapped onto phylogenetic trees.** The carbon content per amino acid side chain and GC1-, GC2-content of the whole genomes are visualized in heatmaps. The branches leading to viral homologs are highlighted in red or green to depict the increase and decrease, respectively, in number of carbons in the viral homolog in comparison to the number of carbons in the ancestral state (located on the other end of the colored branch). The actual change in number of carbons is shown above the branches. The tree was rooted to correspond to the reference phylogeny (**Figure S2**). Scale bar, number of substitutions per site. GC content, in percent. Carbon content, in number of carbons per amino acid.

**Figure S4. The number of carbons and number of high-energy phosphates in proteins encoded by all protein-coding genes in 212 genomes, highly expressed genes, and GTA genes.** Boxplots represent median values that are bounded by the first and third quartiles. Whiskers show the values that lie in the range of 1.5*interquartile rule. Dots outside of the whiskers are outliers.

**Figure S5. Change in relative carbon utilization during the evolutionary histories of GTA genes.** Each tree represents the phylogeny of a GTA gene. The branches are colored to show the dynamics of change in carbon utilization along the branches, which was inferred using the ancestral state reconstruction (see **Materials and Methods** for details). The number in parentheses next to the protein name shows the protein length in the *Rhodobacter capsulatus* GTA. Relative carbon utilization is the ratio of the average number of carbons per amino acid in a GTA gene and in the 26 single-copy genes. Scale bar, number of substitutions per site.

**Figure S6. The relative abundance of amino acids at sites that are inferred to be under the positive selection and contribute to the decrease in the carbon utilization in *Sphingomonadales*.** For each GTA protein, the site number corresponds to the position of the site in the multiple sequence alignment. For each site, the height of an amino is proportional to its frequency in the site. Each amino acid is color-coded by the number of carbons in the side chain of the amino acid (see color legend). For each protein, the lower and upper panels correspond to the amino acid abundances in *Sphingomonadales* and in other three orders. Sites that reduce carbon utilization by more than 3 atoms are outlined with a red rectangle. Arrows indicate sites that were discussed in **Results**.

## Supplementary Table Captions

**Table S1.** The number of detected GTA genes in 212 alphaproteobacterial genomes and viruses from the RefSeq database.

**Table S2.** Functional annotations of 26 single-copy genes that were used to calculate the normalized carbon utilization value.

**Table S3.** The likelihood ratio test for the branch site A model.

**Table S4.** The average number of carbons per amino acid side chain in *Sphigomonadales* and three other orders combined together.

**Table S5.** Contribution of sites under the positive selection to the carbon utilization in *Sphingomonadales’* GTA genes.

## Supplementary Movie Caption

**Movie S1. Visualization of the inferred positively selected sites in *Sphingomonadales* on a 3D structural model of the T5 bacteriophage major capsid protein.** The hexamer structure is shown in green as a cartoon of secondary structure elements. Atoms of the amino acids that are inferred to be under the positive selection in *Sphingomonadales* are highlighted as pink spheres. The structure is rotated to facilitate visualization of the position of the positively selected amino acids.

