## Supplementary figures and images for "Selection for reducing energy cost of protein production drives the GC content and amino acid composition bias in gene transfer agents"

### Supplemental Figure S1

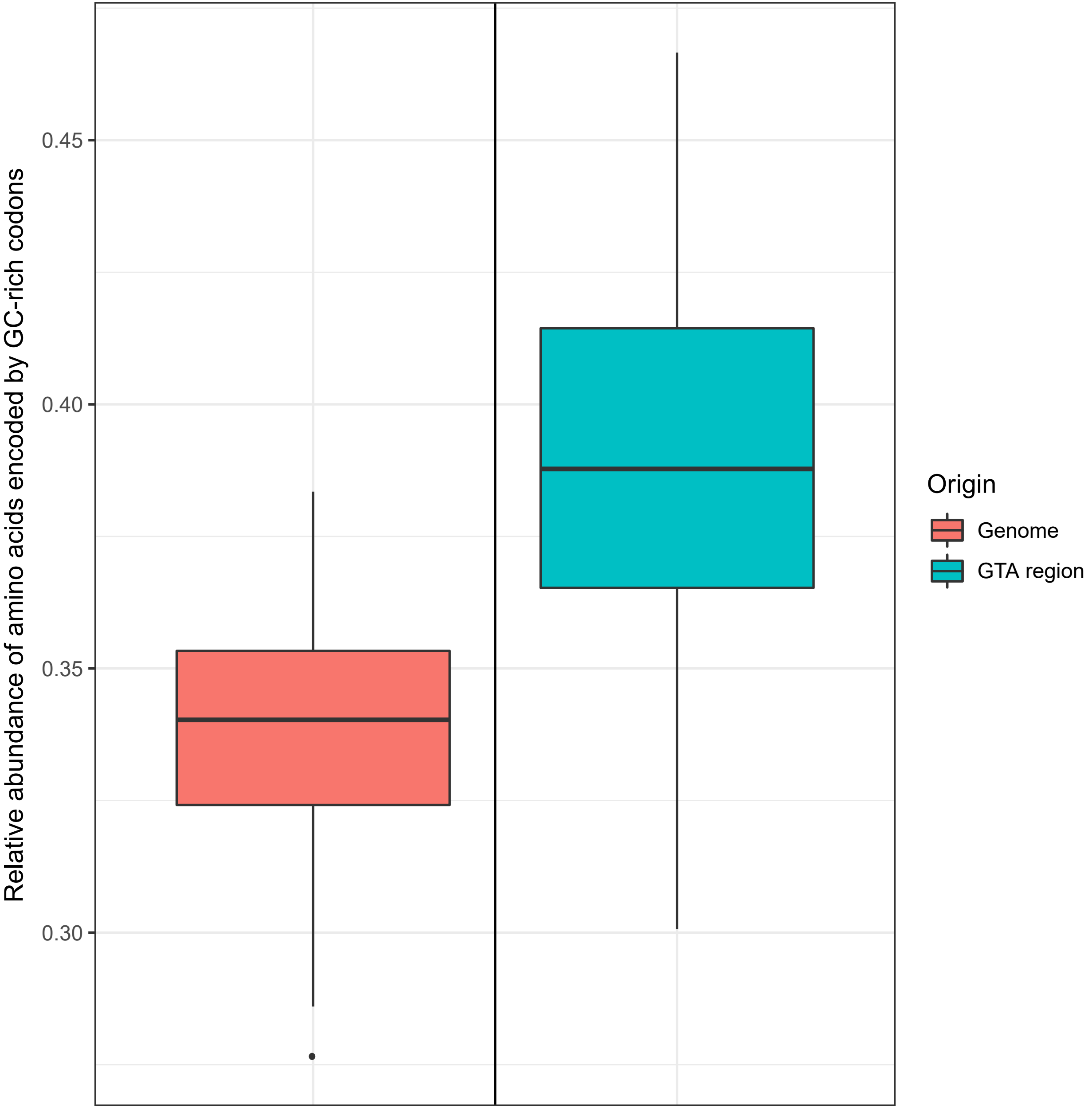

### Supplemental Figure S2

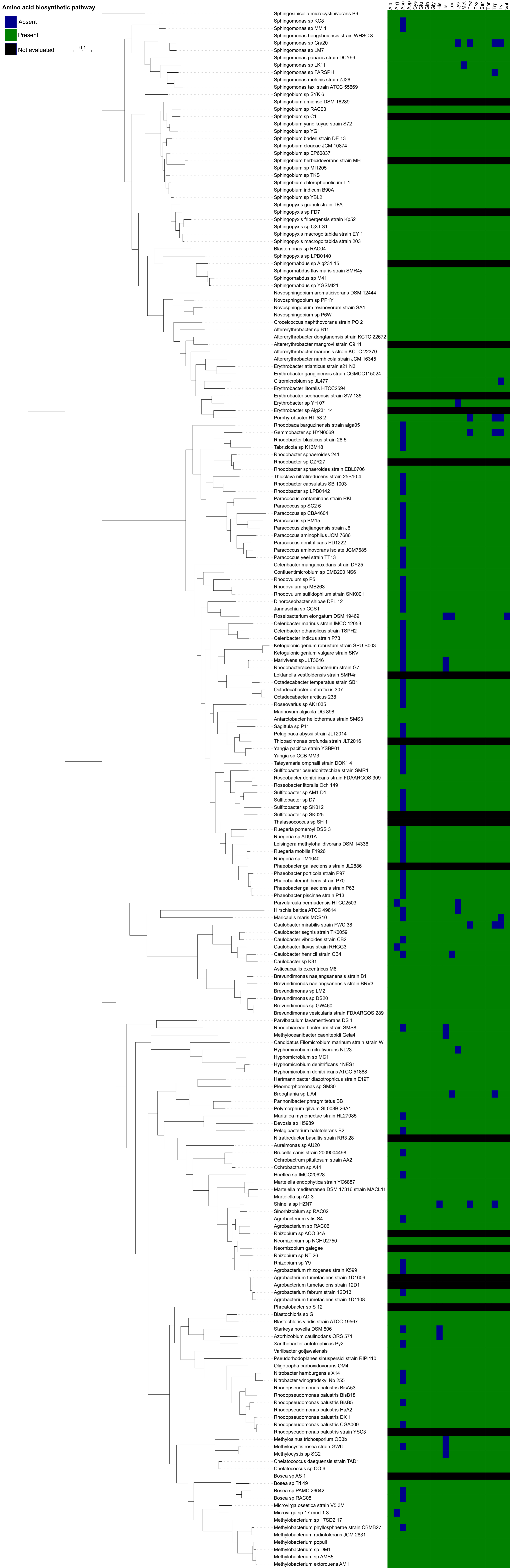

### Supplemental Figure S3

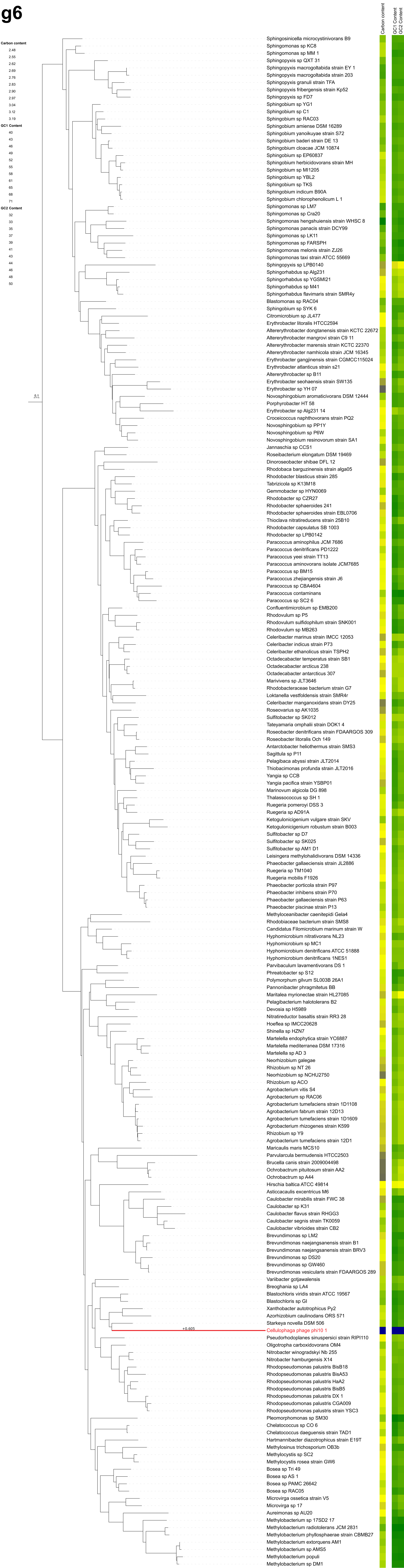

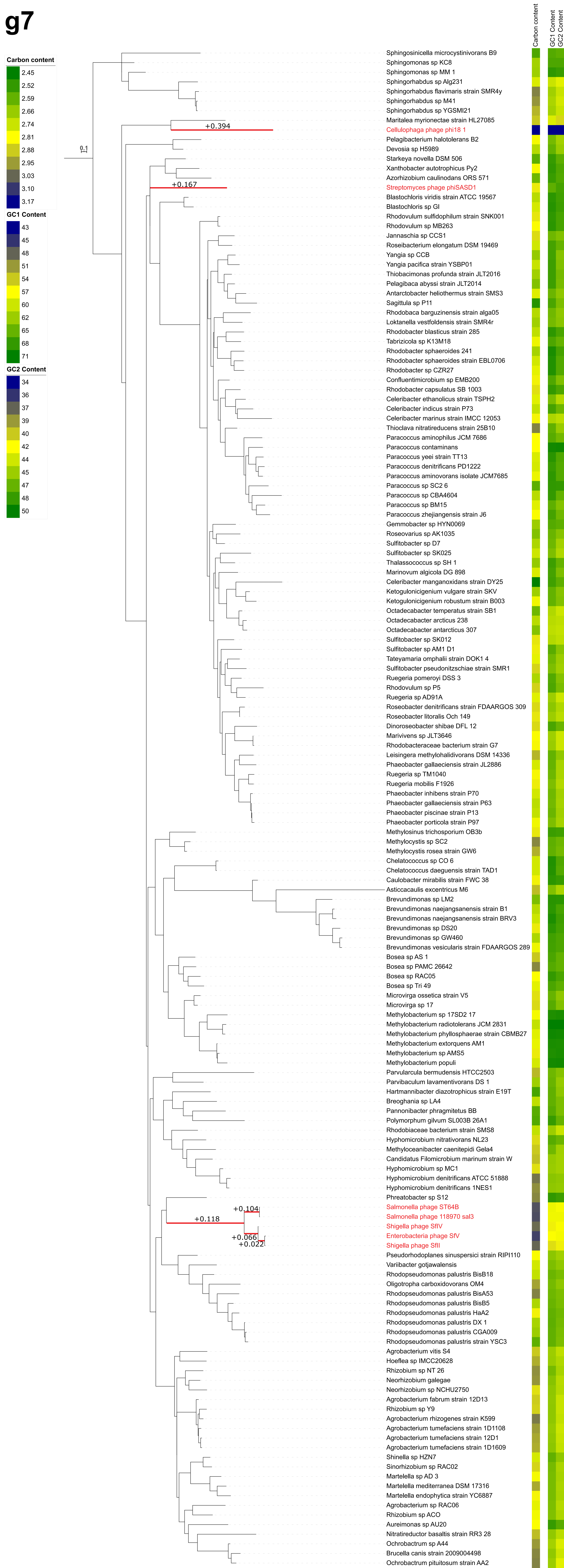

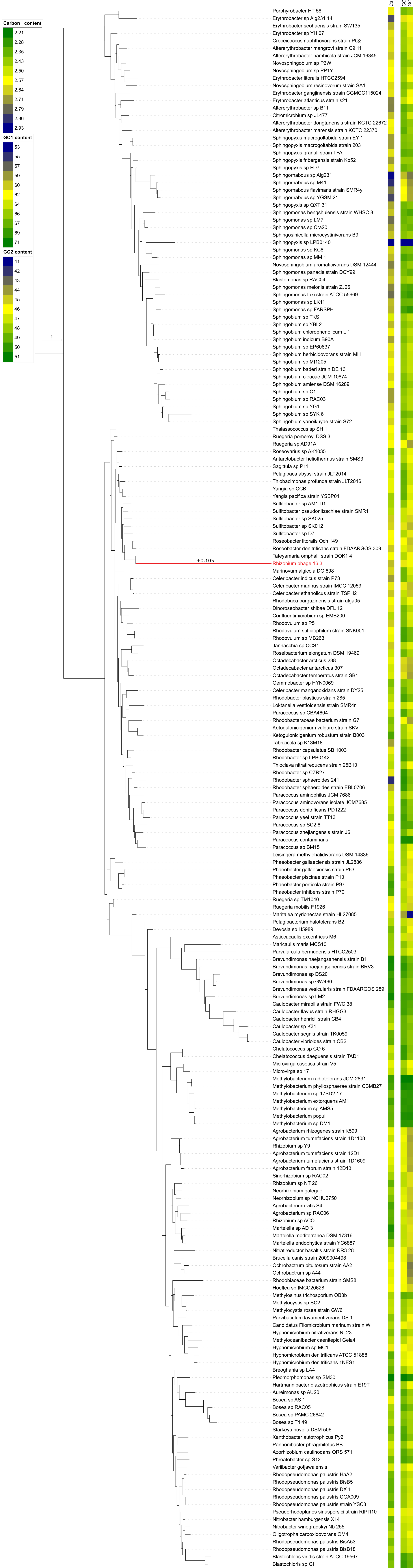

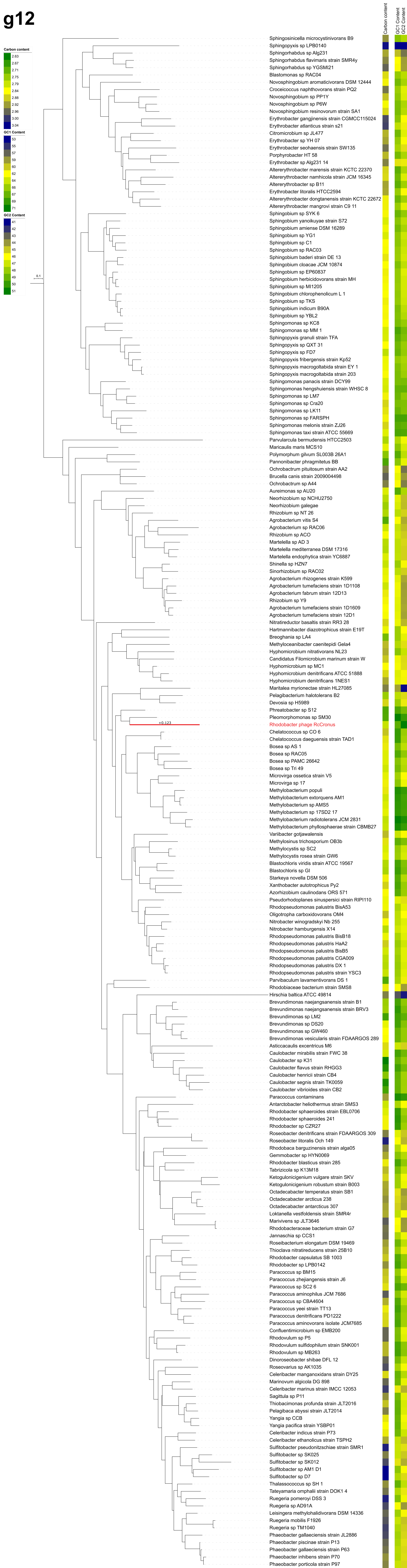

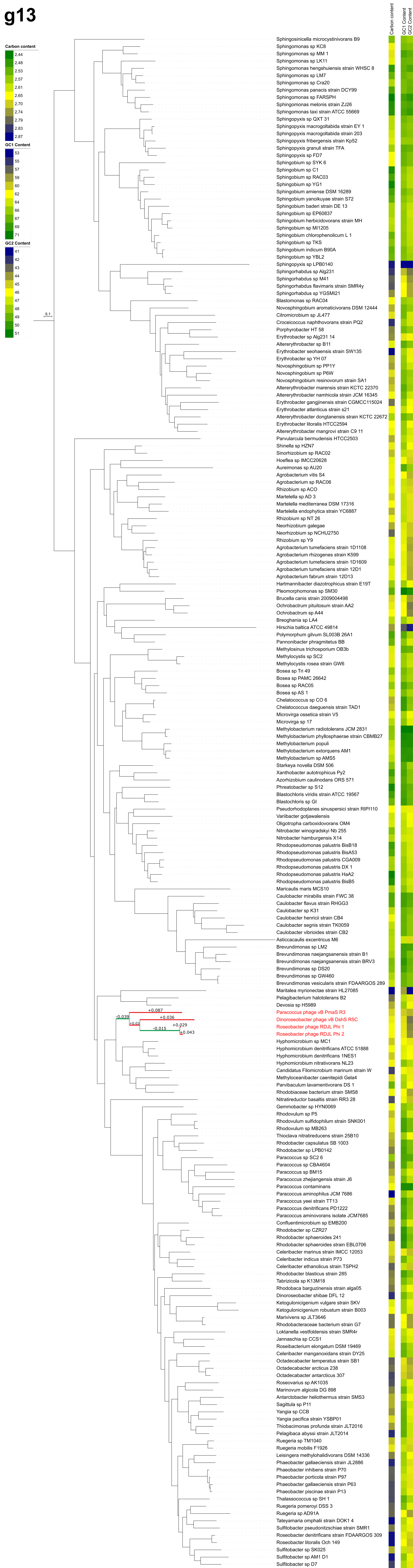

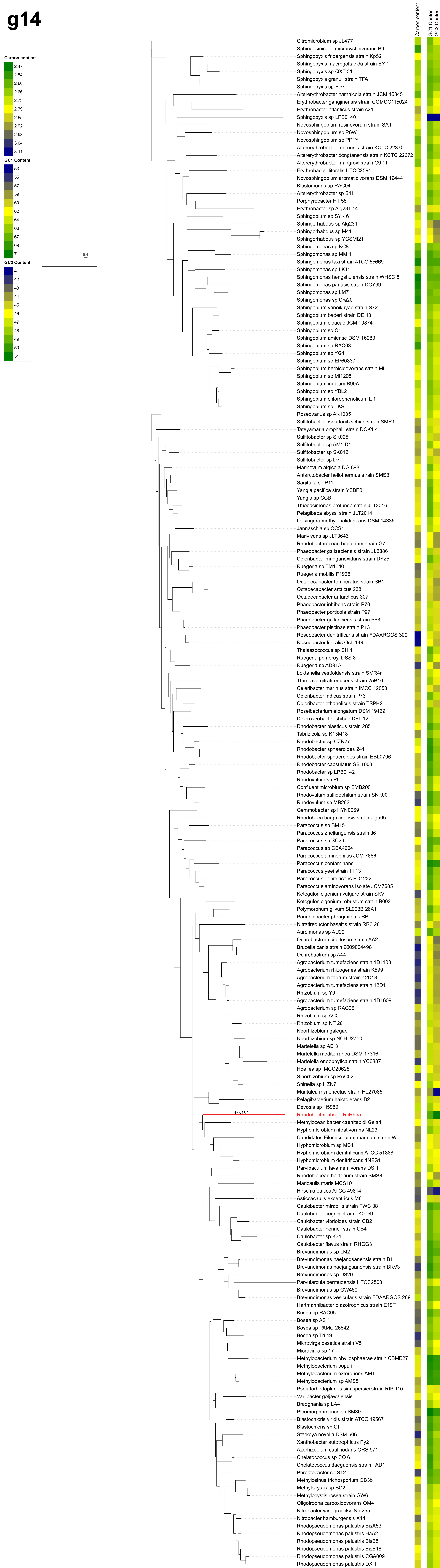

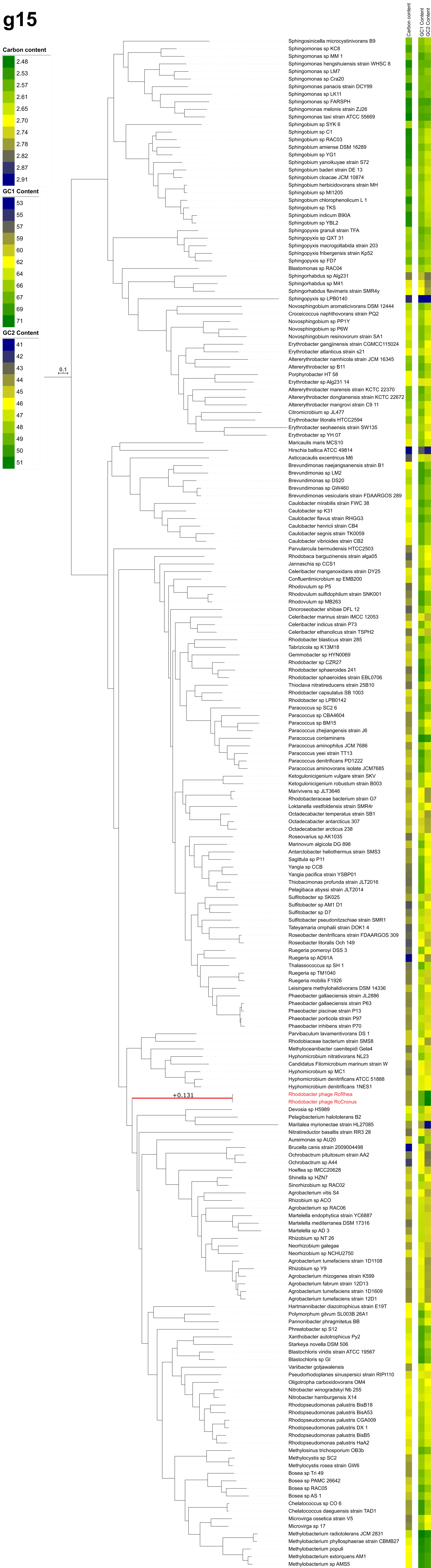

### Supplemental Figure S4

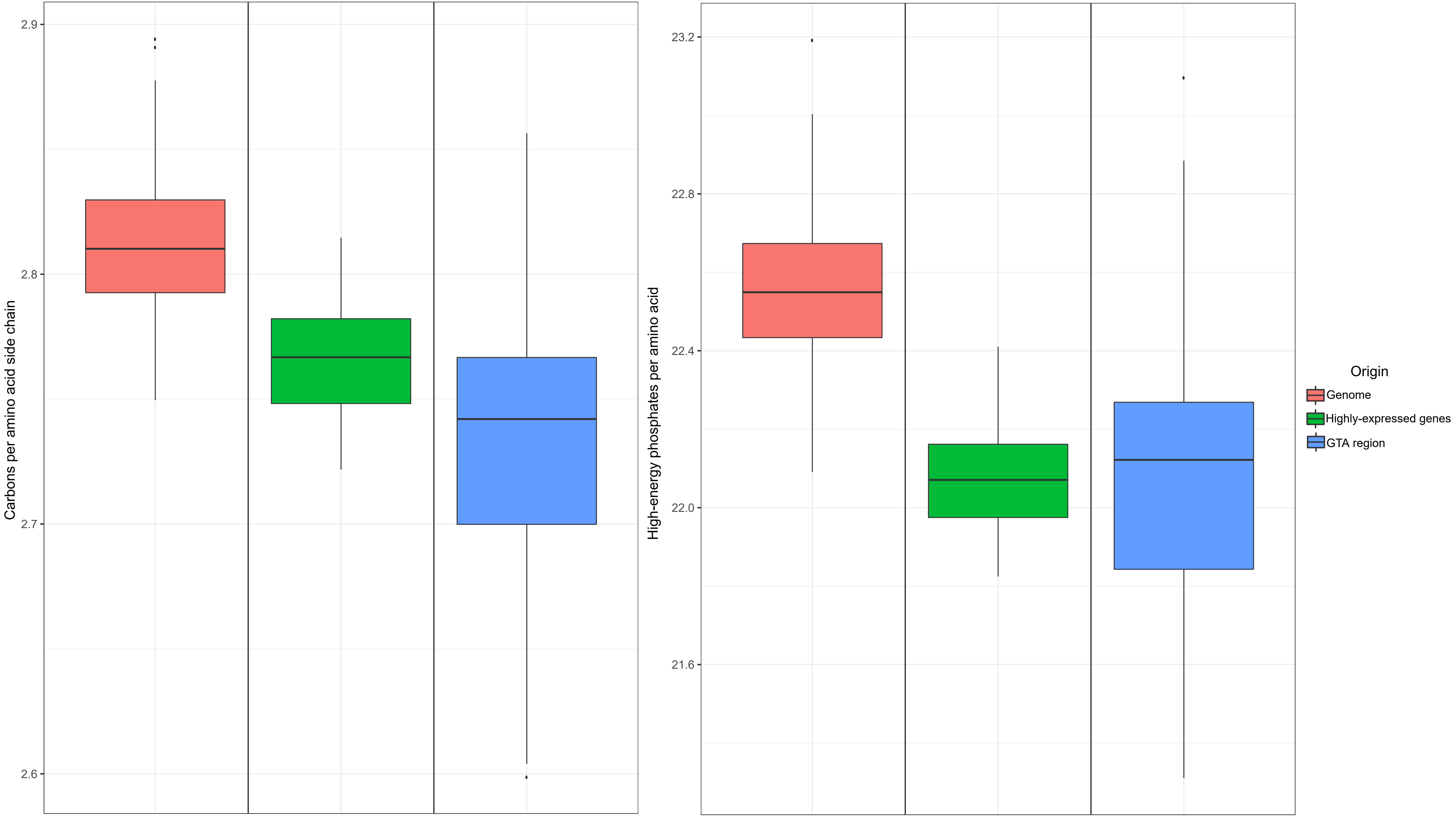

### Supplemental Figure S6

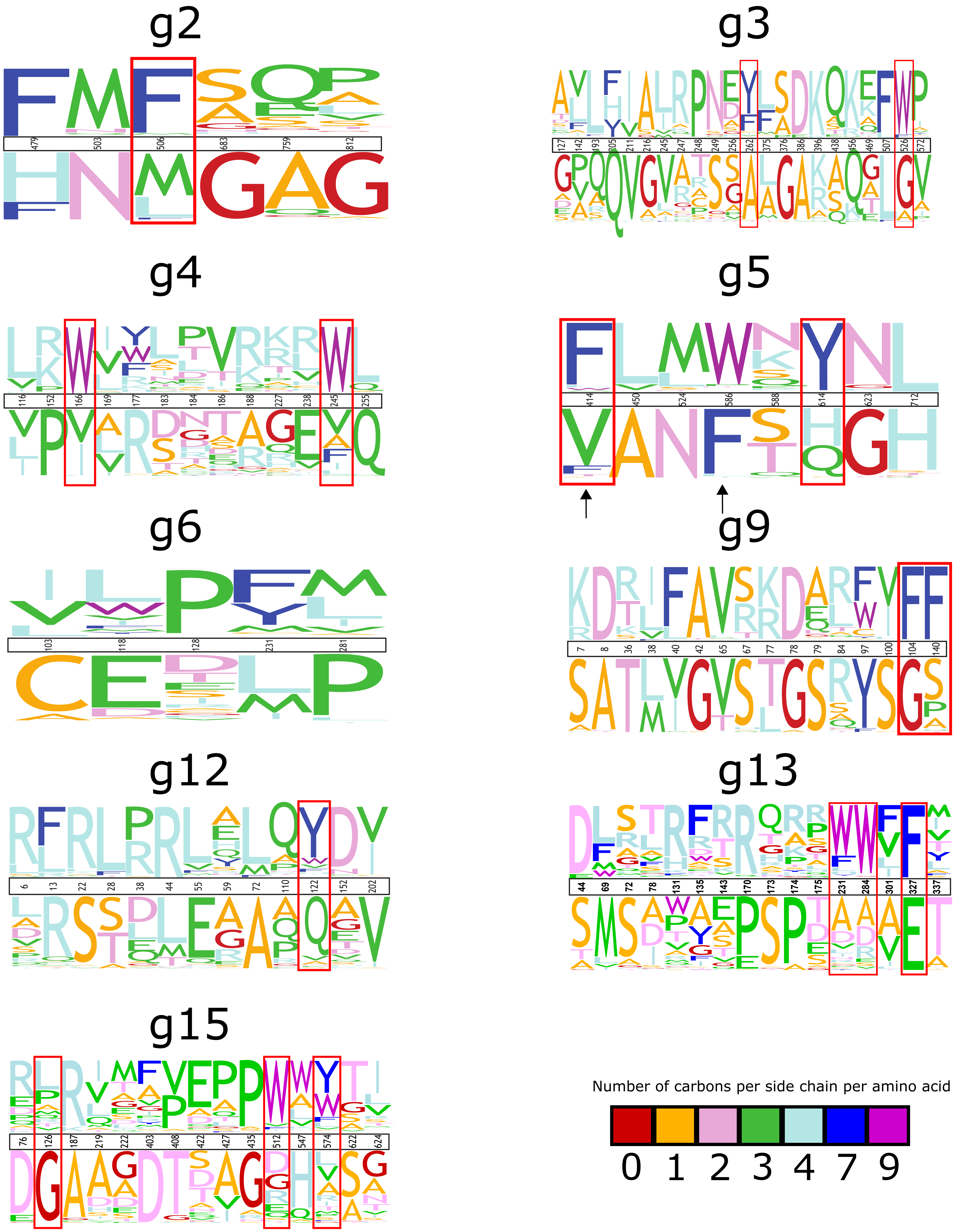
