## Supplemental Figure S5 for "Selection for reducing energy cost of protein production drives the GC content and amino acid composition bias in gene transfer agents"

g2
(448 aa)

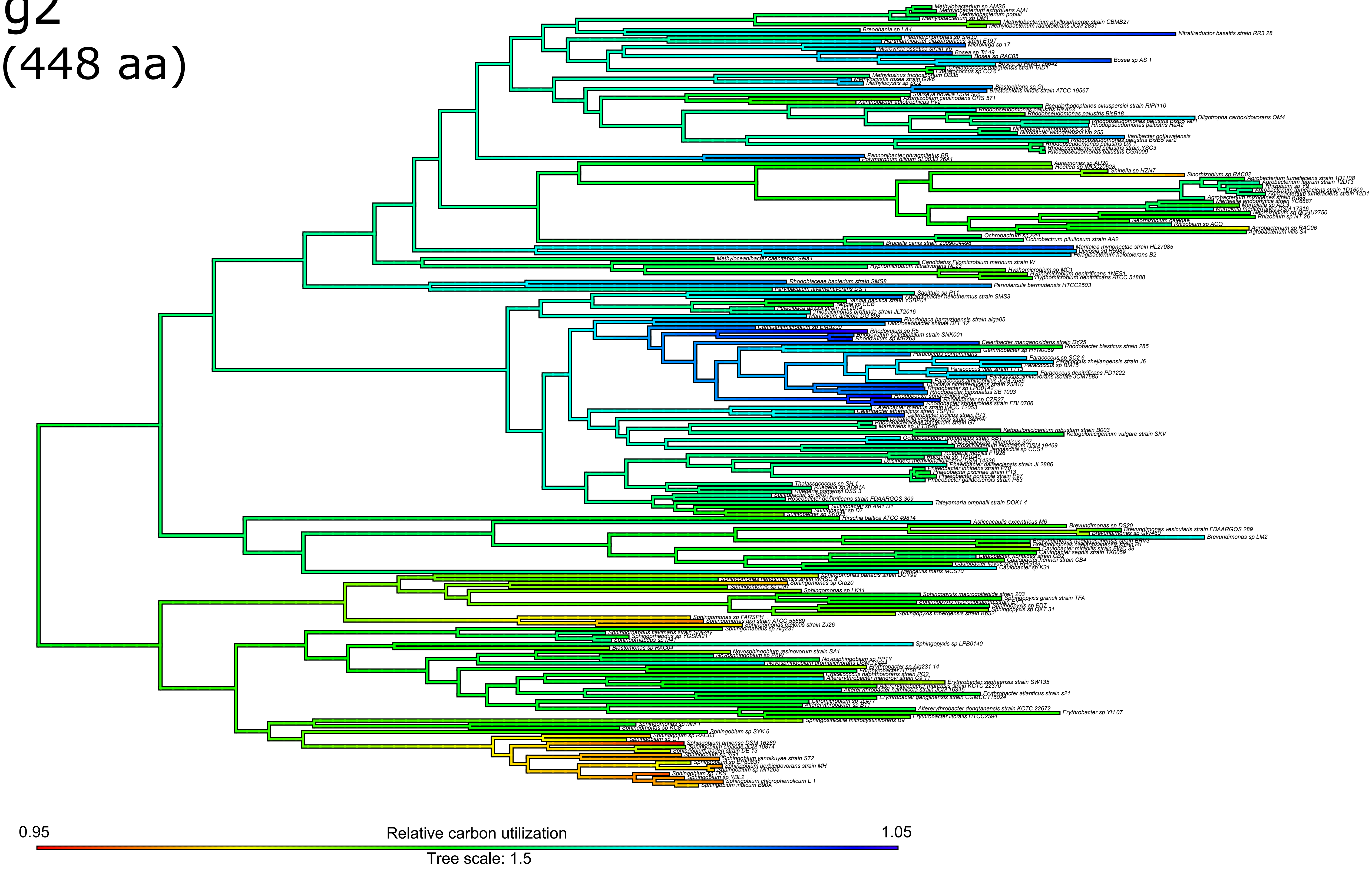

g3
(396 aa)

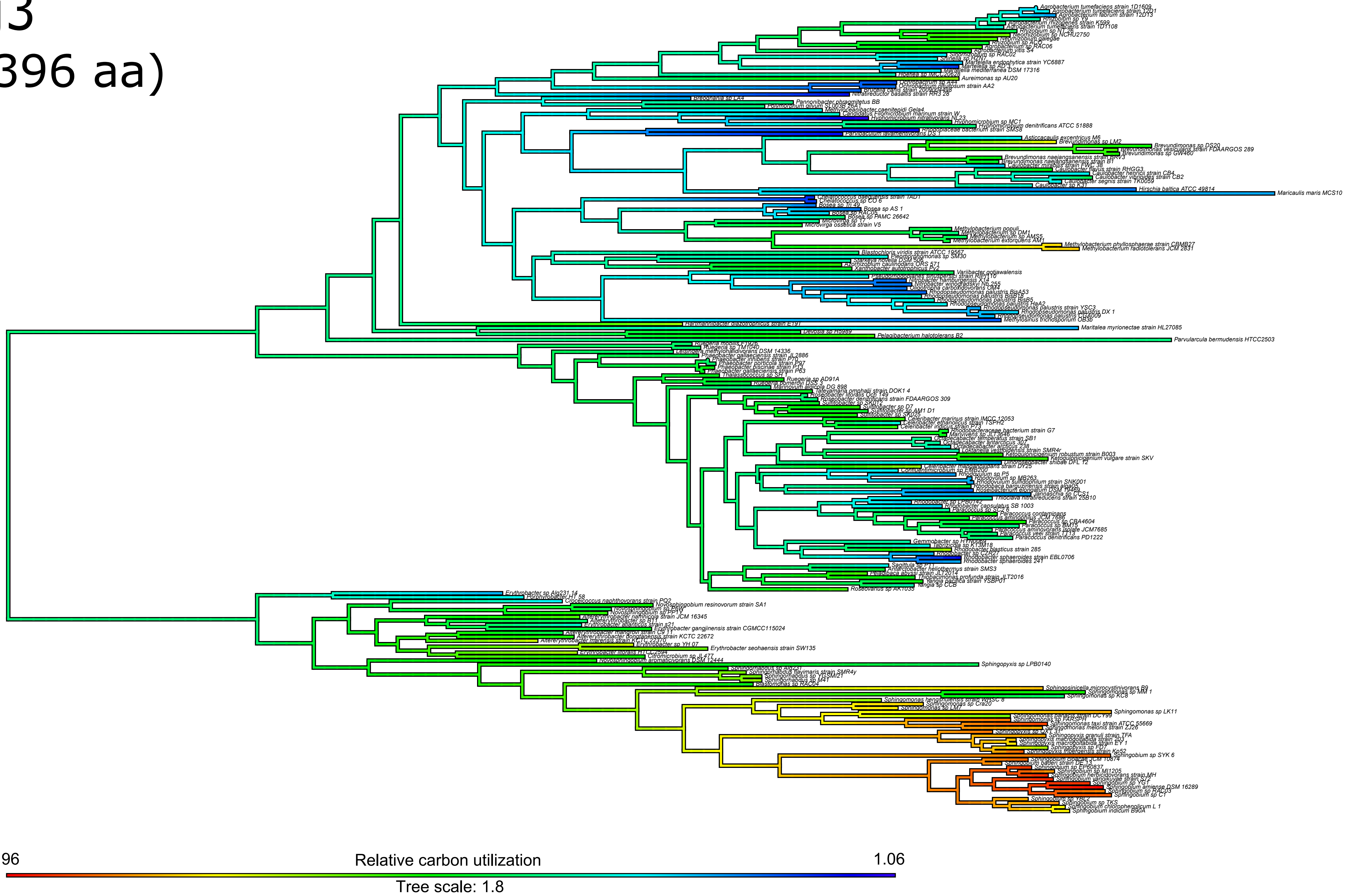

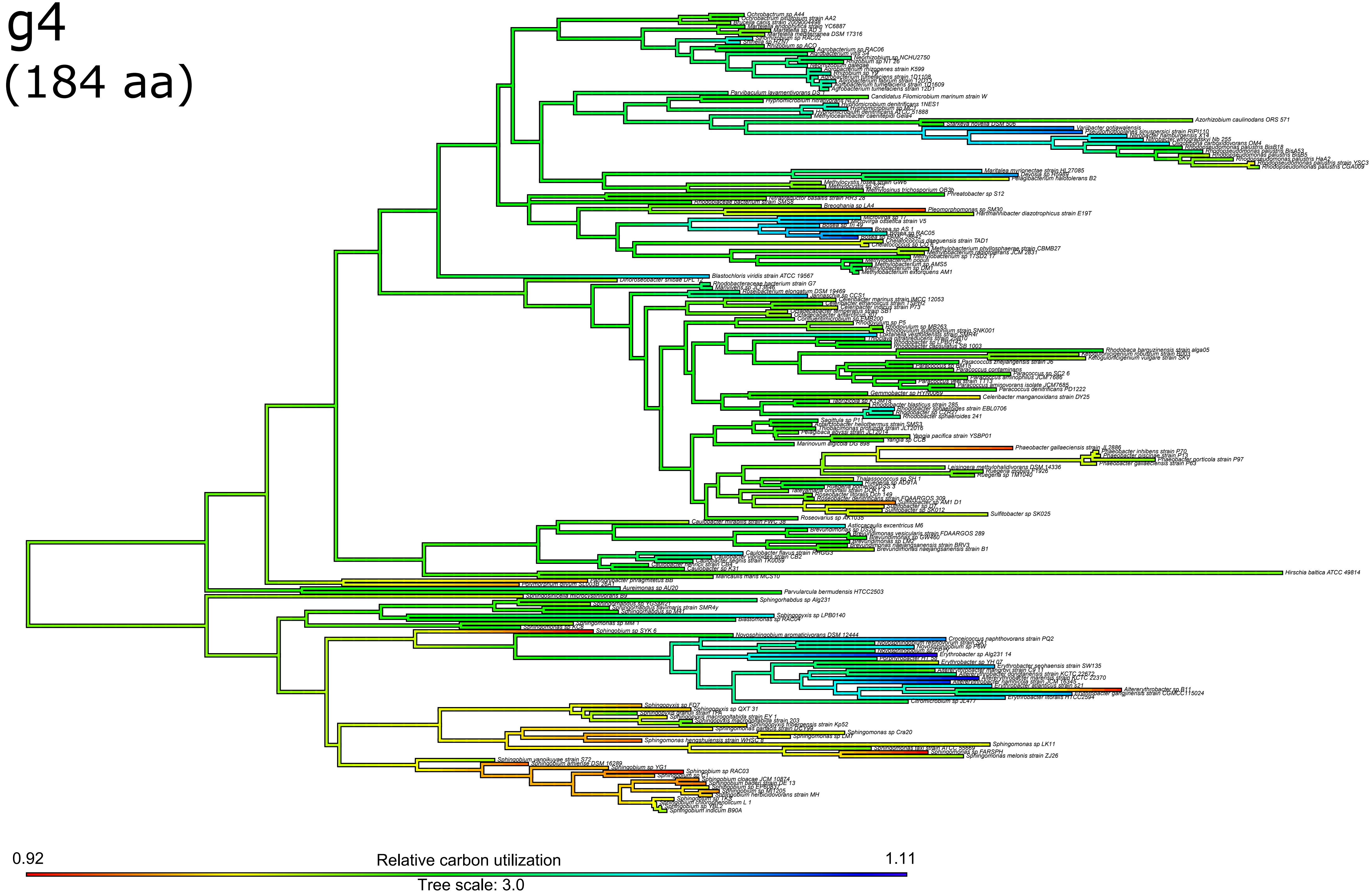
g4
(184 aa)
0.92
Relative carbon utilization
1.11
Tree scale: 3.0

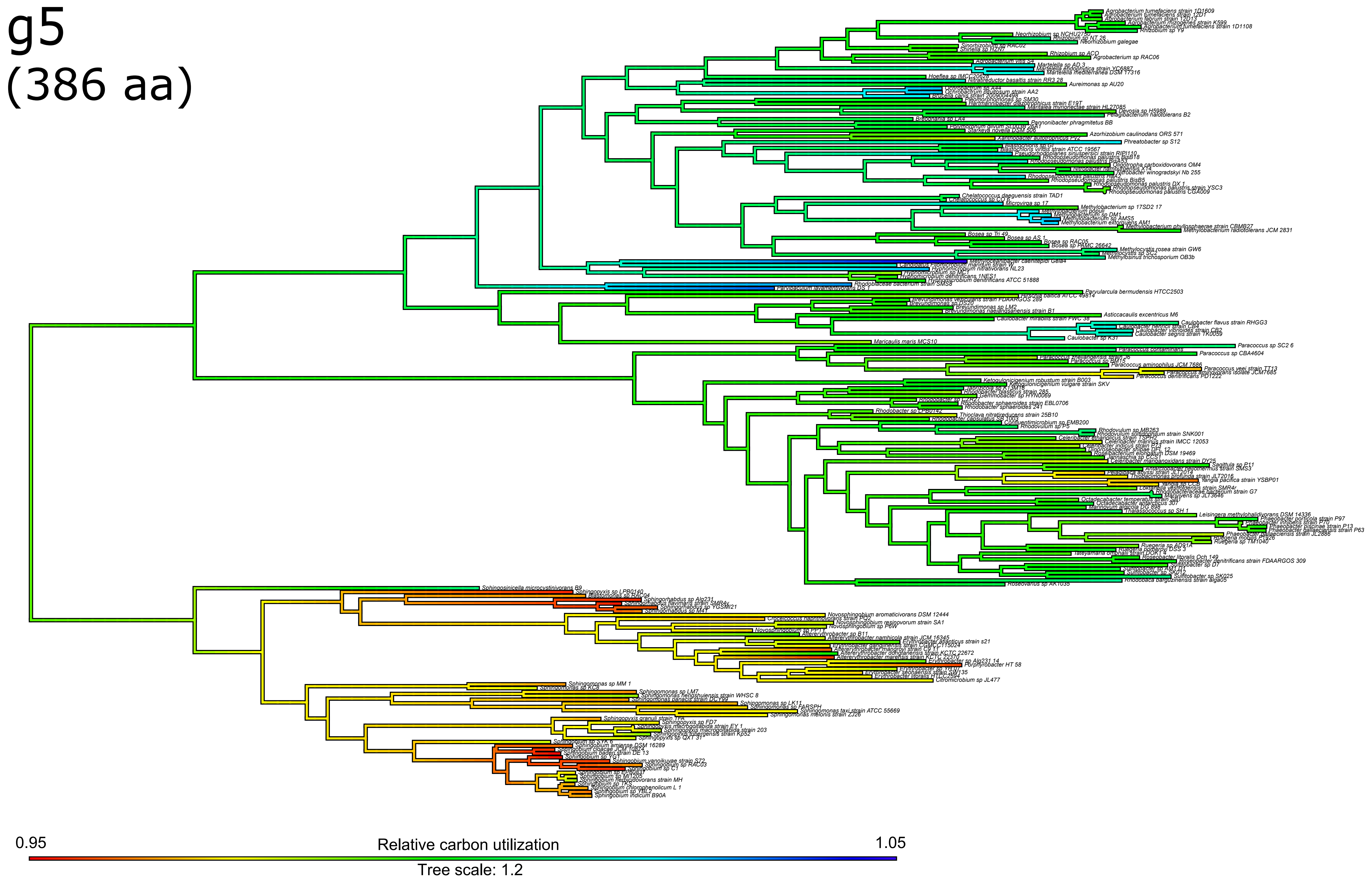
g5
(386 aa)
0.95
Relative carbon utilization
1.05
Tree scale: 1.2

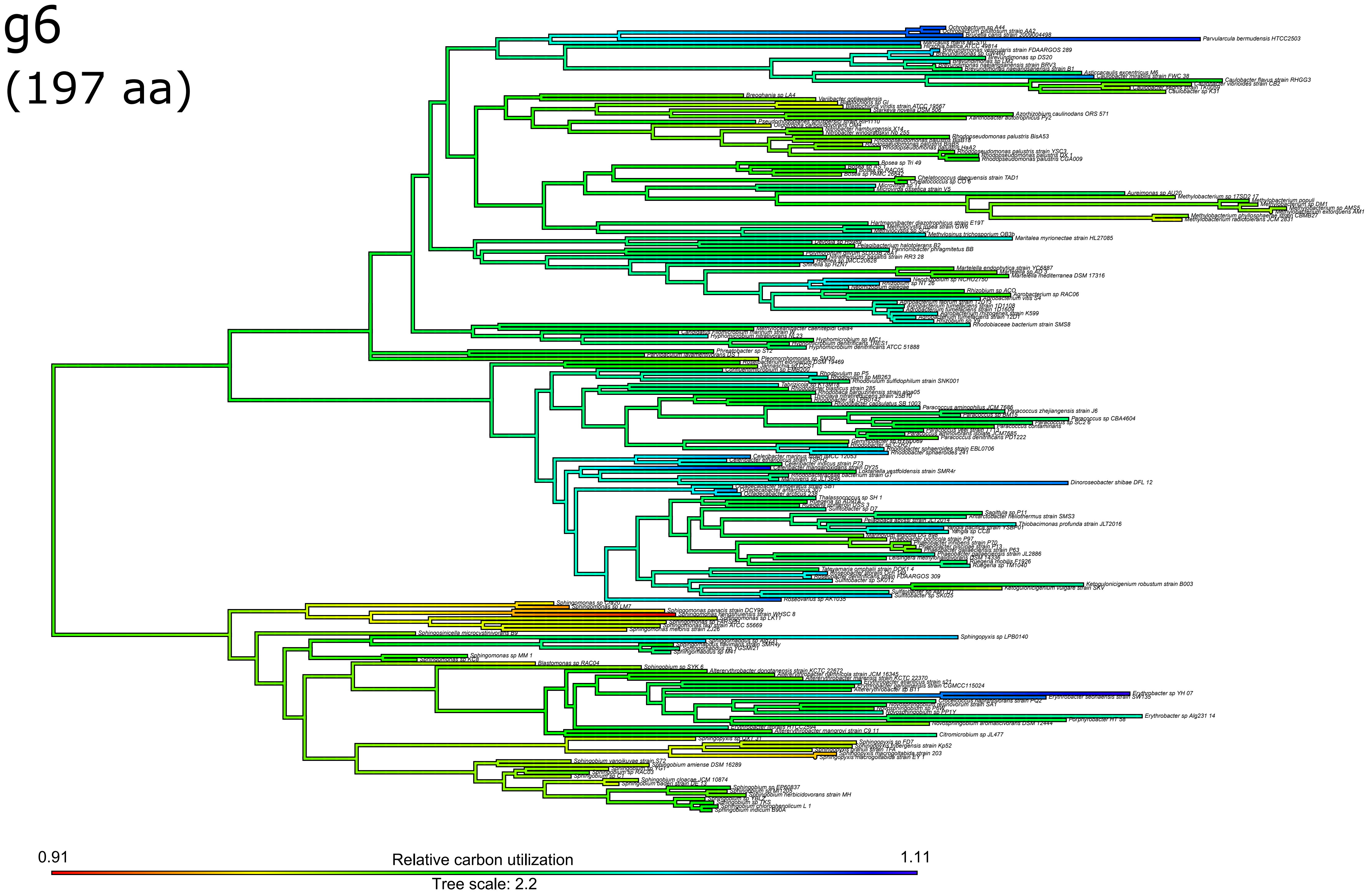
g6
(197 aa)
0.91
Relative carbon utilization
1.11
Tree scale: 2.2

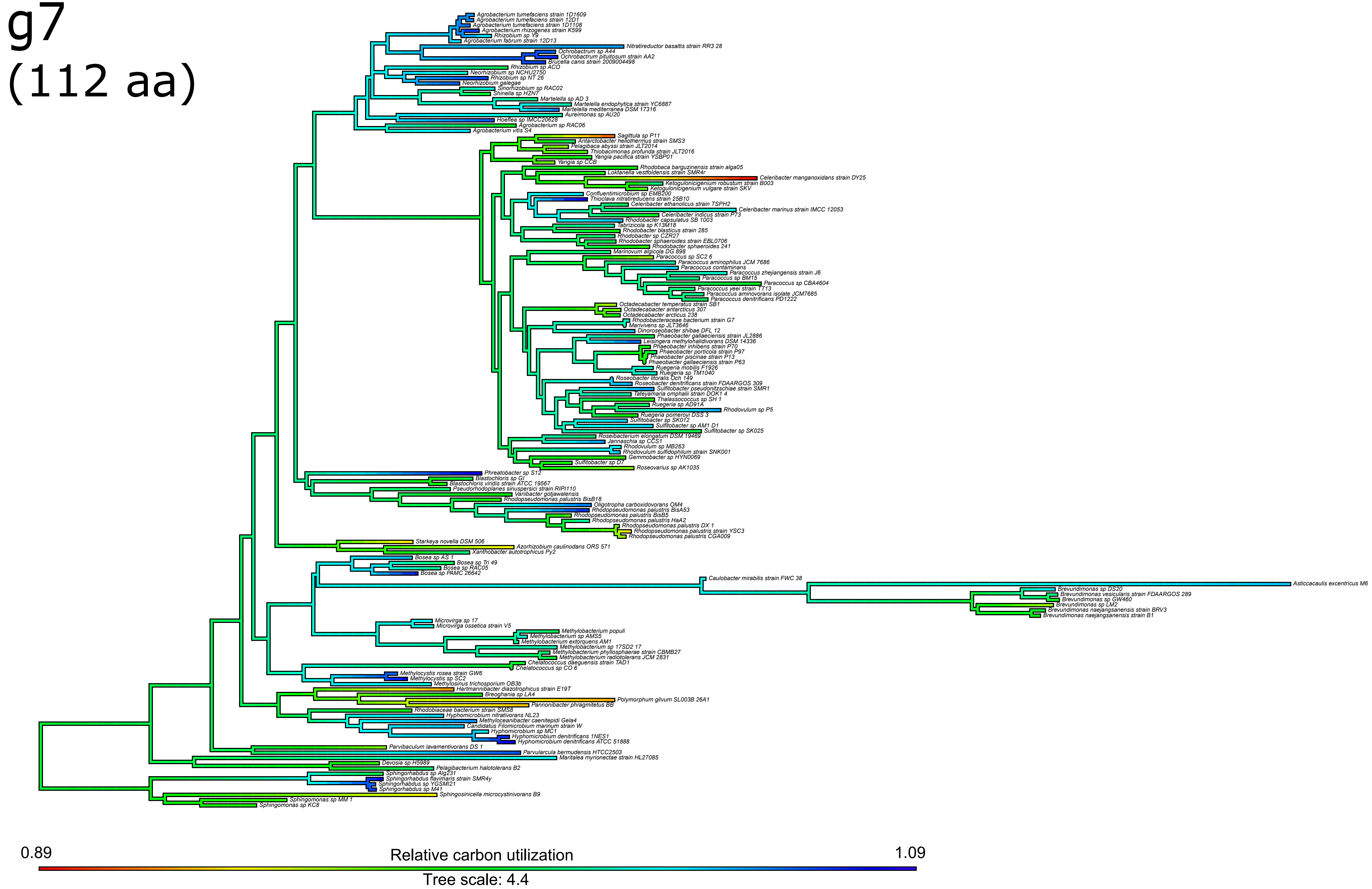
g7
(112 aa)
Agrobacterium tumefaciens strain 1D1609
Agrobacterium tumefaciens strain 12D1
Agrobacterium tumefaciens strain 1D1108
Agrobacterium rhizogenes strain K599
Rhizobium sp Y9
Agrobacterium fabrum strain 12D13
Nitratireductor basaltis strain RR3 28
Ochrobactrum sp A44
Ochrobactrum pituitosum strain AA2
Brucella canis strain 2009004498
Rhizobium sp ACO
Neorhizobium sp NCHU2750
Rhizobium sp NT 26
Neorhizobium galegae
Sinorhizobium sp RAC02
Shinella sp HZN7
Martelella sp AD 3
Martelella endophytica strain YC6887
Martelella mediterranea DSM 17316
Aureimonas sp AU20
Hoeflea sp IMCC20628
Agrobacterium sp RAC06
Agrobacterium vitis S4
Sagittula sp P11
Antarctobacter heliothermus strain SMS3
Pelagibaca abyssi strain JLT2014
Thiobacimonas profunda strain JLT2016
Yangia pacifica strain YSBP01
Yangia sp CCB
Rhodobaca barguzinensis strain alga05
Loktanella vestfoldensis strain SMR4r
Celeribacter manganoxidans strain DY25
Ketogulonicigenium robustum strain B003
Ketogulonicigenium vulgare strain SKV
Confluentimicrobium sp EMB200
Thioclava nitratireducens strain 25B10
Celeribacter ethanolicus strain TSPH2
Celeribacter marinus strain IMCC 12053
Celeribacter indicus strain P73
Rhodobacter capsulatus SB 1003
Tabrizicola sp K13M18
Rhodobacter blasticus strain 285
Rhodobacter sp CZR27
Rhodobacter sphaeroides strain EBL0706
Rhodobacter sphaeroides 241
Marinovum algicola DG 898
Paracoccus sp SC2 6
Paracoccus aminophilus JCM 7686
Paracoccus contaminans
Paracoccus zhejiangensis strain J6
Paracoccus sp BM15
Paracoccus sp CBA4604
Paracoccus yeei strain TT13
Paracoccus aminovorans isolate JCM7685
Paracoccus denitrificans PD1222
Octadecabacter temperatus strain SB1
Octadecabacter antarcticus 307
Octadecabacter arcticus 238
Rhodobacteraceae bacterium strain G7
Marivivens sp JLT3646
Dinoroseobacter shibae DFL 12
Phaeobacter gallaeciensis strain JL2886
Leisingera methylohalidivorans DSM 14336
Phaeobacter inhibens strain P70
Phaeobacter porticola strain P97
Phaeobacter piscinae strain P13
Phaeobacter gallaeciensis strain P63
Ruegeria mobilis F1926
Ruegeria sp TM1040
Roseobacter litoralis Och 149
Roseobacter denitrificans strain FDAARGOS 309
Sulfitobacter pseudonitzschiae strain SMR1
Tateyamaria omphalii strain DOK1 4
Thalassococcus sp SH 1
Ruegeria sp AD91A
Rhodovulum sp P5
Ruegeria pomeroyi DSS 3
Sulfitobacter sp SK012
Sulfitobacter sp AM1 D1
Sulfitobacter sp SK025
Roseibacterium elongatum DSM 19469
Jannaschia sp CCS1
Rhodovulum sp MB263
Rhodovulum sulfidophilum strain SNK001
Gemmobacter sp HYN0069
Sulfitobacter sp D7
Roseovarius sp AK1035
Phreatobacter sp S12
Blastochloris sp GI
Blastochloris viridis strain ATCC 19567
Pseudorhodoplanes sinuspersici strain RIPI110
Variibacter gotjawalensis
Rhodopseudomonas palustris BisB18
Oligotropha carboxidovorans OM4
Rhodopseudomonas palustris BisA53
Rhodopseudomonas palustris BisB5
Rhodopseudomonas palustris HaA2
Rhodopseudomonas palustris DX 1
Rhodopseudomonas palustris strain YSC3
Rhodopseudomonas palustris CGA009
Starkeya novella DSM 506
Azorhizobium caulinodans ORS 571
Xanthobacter autotrophicus Py2
Bosea sp AS 1
Bosea sp Tri 49
Bosea sp RAC05
Bosea sp PAMC 26642
Caulobacter mirabilis strain FWC 38
Asticcacaulis excentricus M6
Brevundimonas sp DS20
Brevundimonas vesicularis strain FDAARGOS 289
Brevundimonas sp GW460
Brevundimonas sp LM2
Brevundimonas naejangsanensis strain BRV3
Brevundimonas naejangsanensis strain B1
Microvirga sp 17
Microvirga ossetica strain V5
Methylobacterium populi
Methylobacterium sp AMS5
Methylobacterium extorquens AM1
Methylobacterium sp 17SD2 17
Methylobacterium phyllosphaerae strain CBMB27
Methylobacterium radiotolerans JCM 2831
Chelatococcus daeguensis strain TAD1
Chelatococcus sp CO 6
Methylocystis rosea strain GW6
Methylocystis sp SC2
Methylosinus trichosporium OB3b
Hartmannibacter diazotrophicus strain E19T
Breoghania sp LA4
Polymorphum gilvum SL003B 26A1
Pannonibacter phragmitetus BB
Rhodobiaceae bacterium strain SMS8
Hyphomicrobium nitrativorans NL23
Methyloceanibacter caenitepidi Gela4
Candidatus Filomicrobium marinum strain W
Hyphomicrobium sp MC1
Hyphomicrobium denitrificans 1NES1
Hyphomicrobium denitrificans ATCC 51888
Parvibaculum lavamentivorans DS 1
Parvularcula bermudensis HTCC2503
Maritalea myrionectae strain HL27085
Devosia sp H5989
Pelagibacterium halotolerans B2
Sphingorhabdus sp Alg231
Sphingorhabdus flavimaris strain SMR4y
Sphingorhabdus sp YGSMI21
Sphingorhabdus sp M41
Sphingosinicella microcystinivorans B9
Sphingomonas sp MM 1
Sphingomonas sp KC8
0.89
Relative carbon utilization
1.09
Tree scale: 4.4

g8
(135 aa)

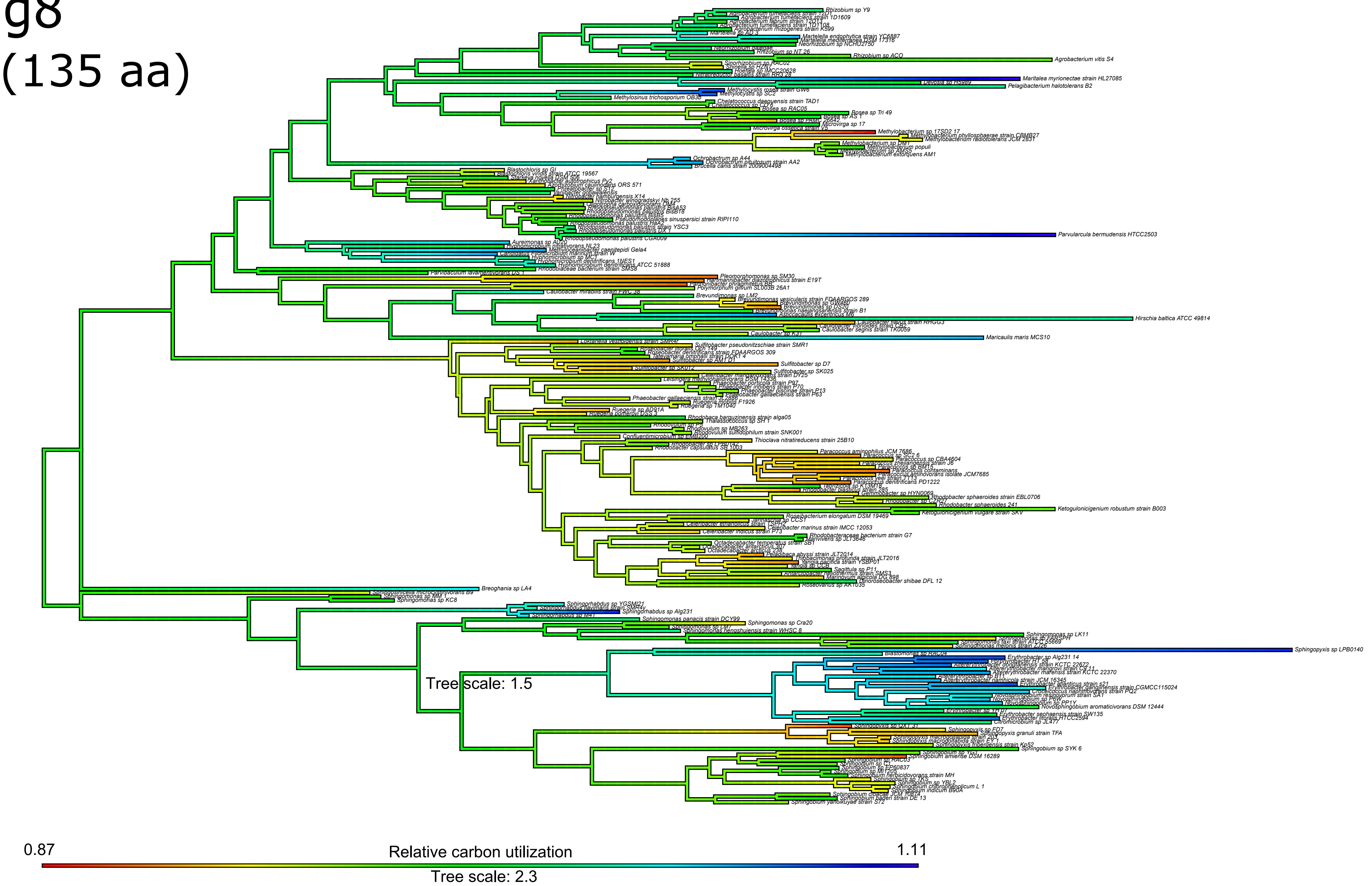
Tree scale: 1.5
0.87
Relative carbon utilization
1.11
Tree scale: 2.3

# g9
(137 aa)

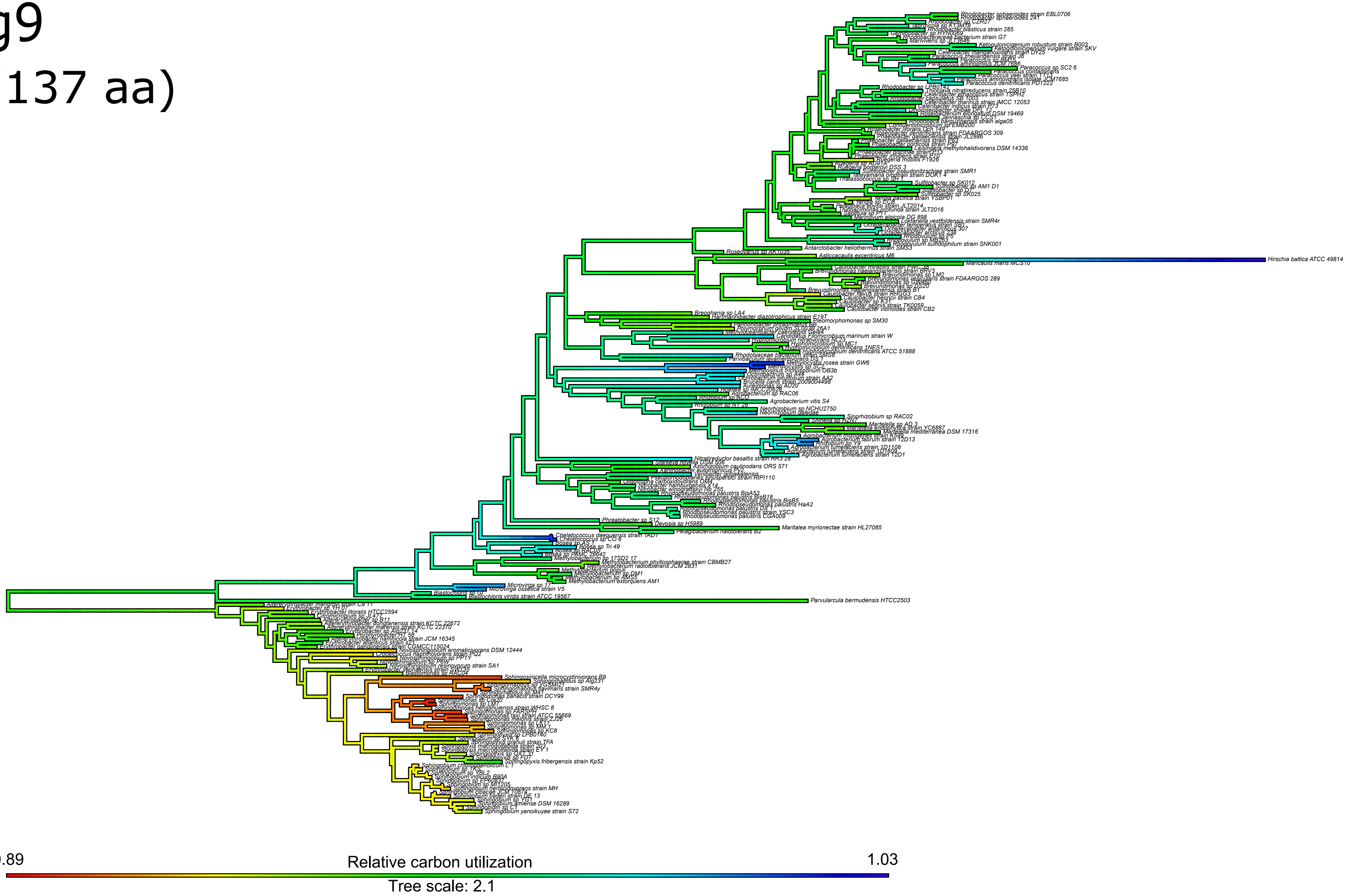
0.89
Relative carbon utilization
1.03
Tree scale: 2.1
Hirschia baltica ATCC 49814
Parvularcula bermudensis HTCC2503

# g10
(108 aa)

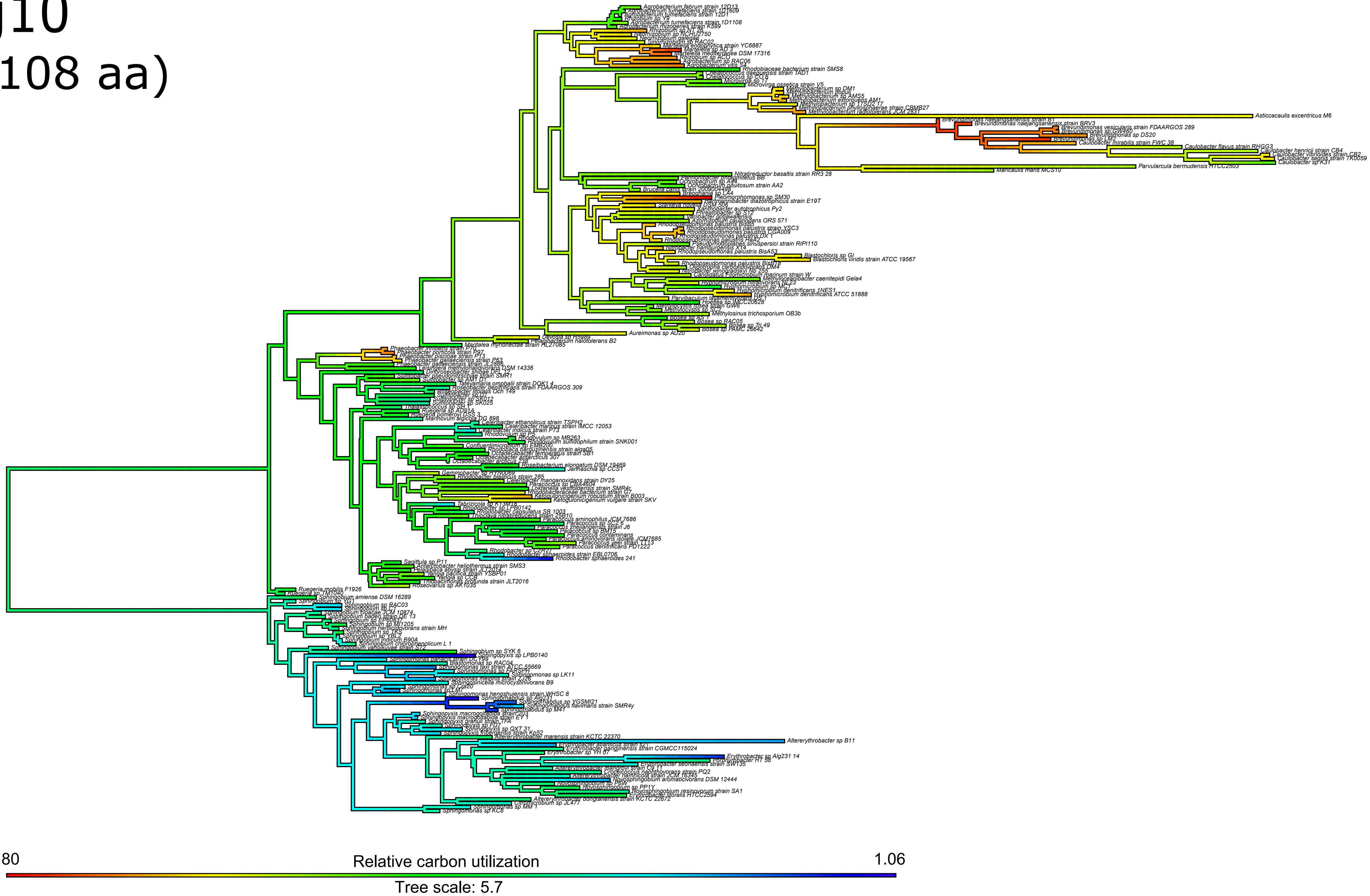

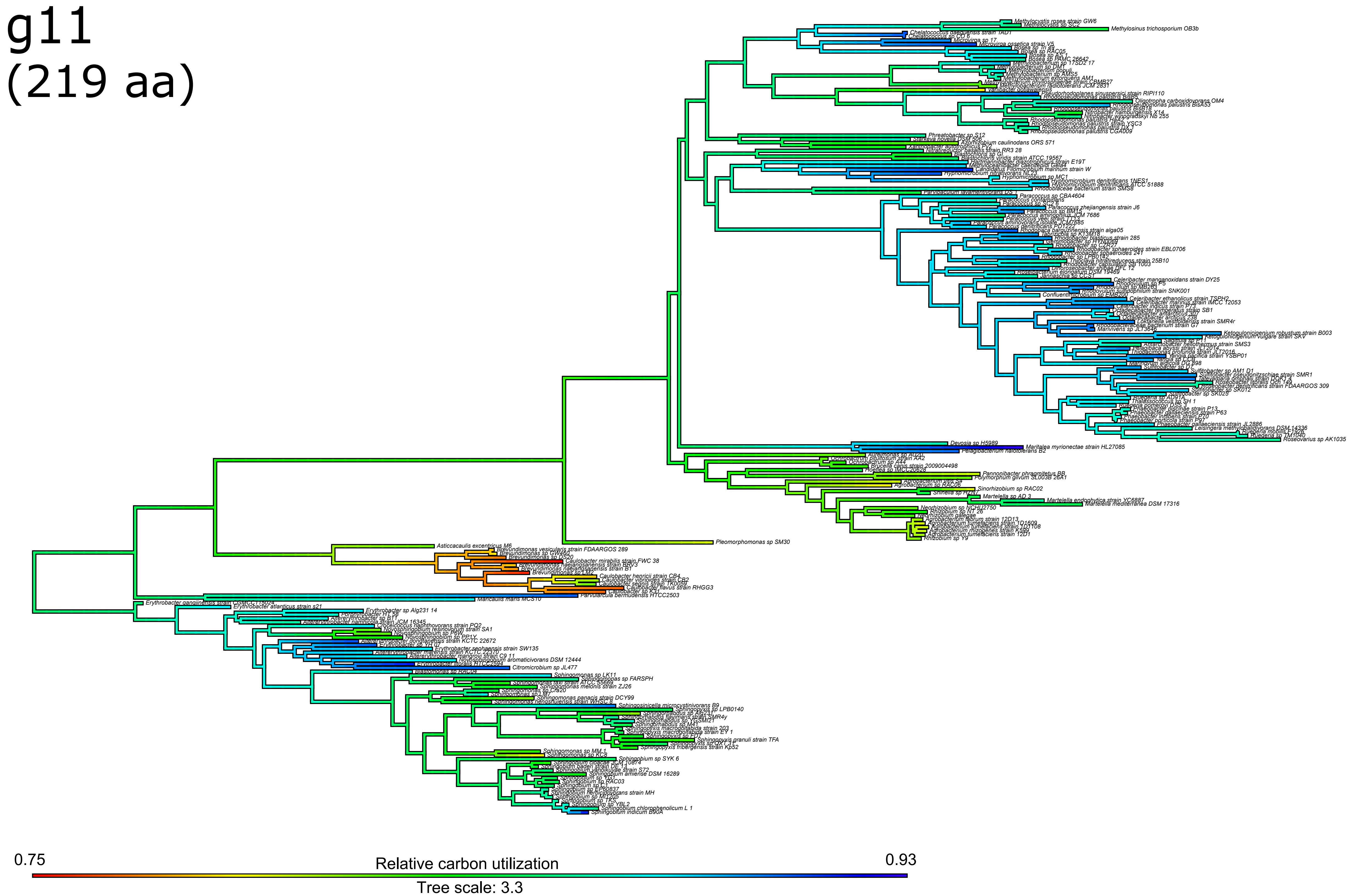
g11
(219 aa)
0.75
Relative carbon utilization
0.93
Tree scale: 3.3

# g12
# (210 aa)

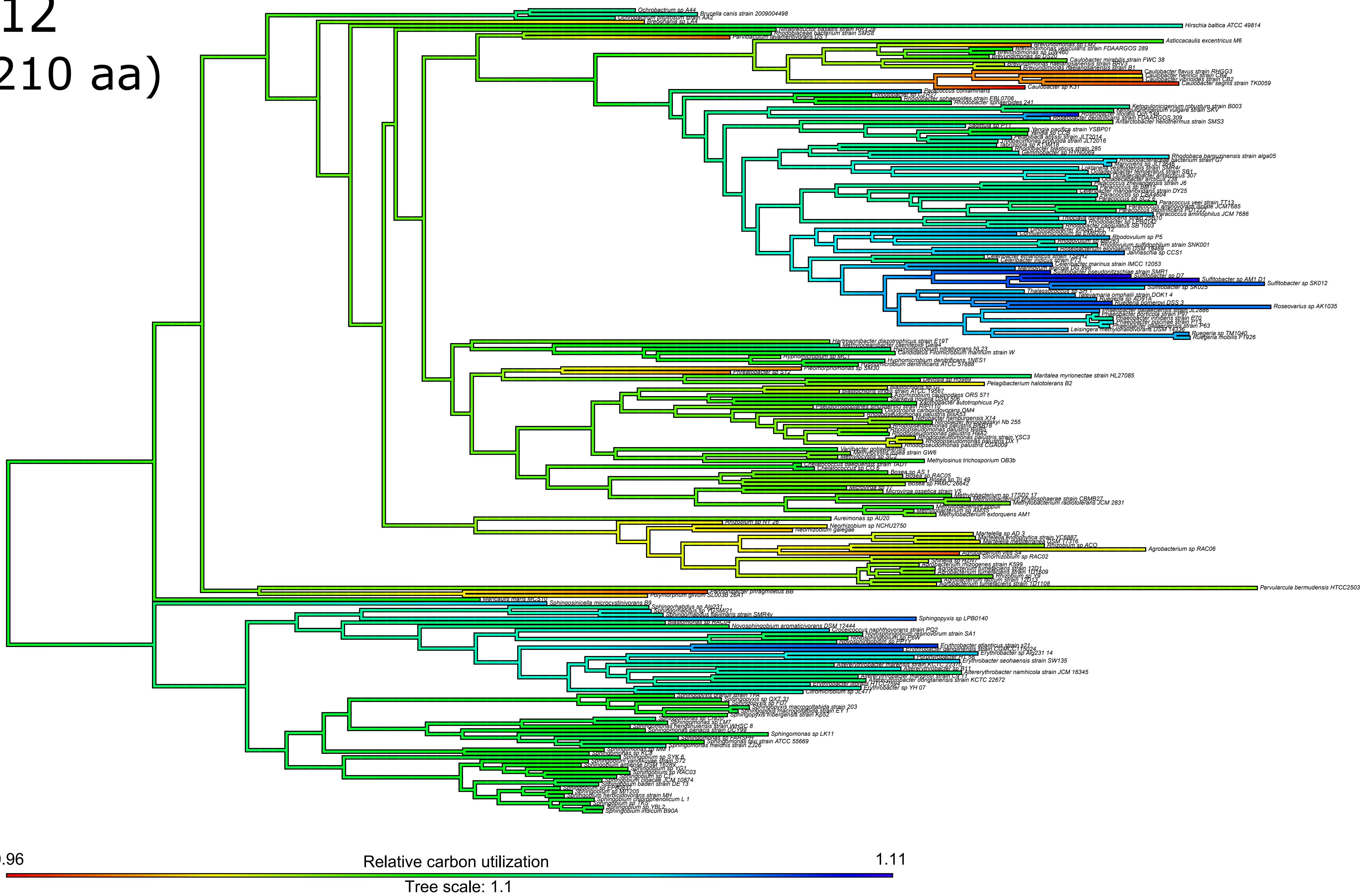

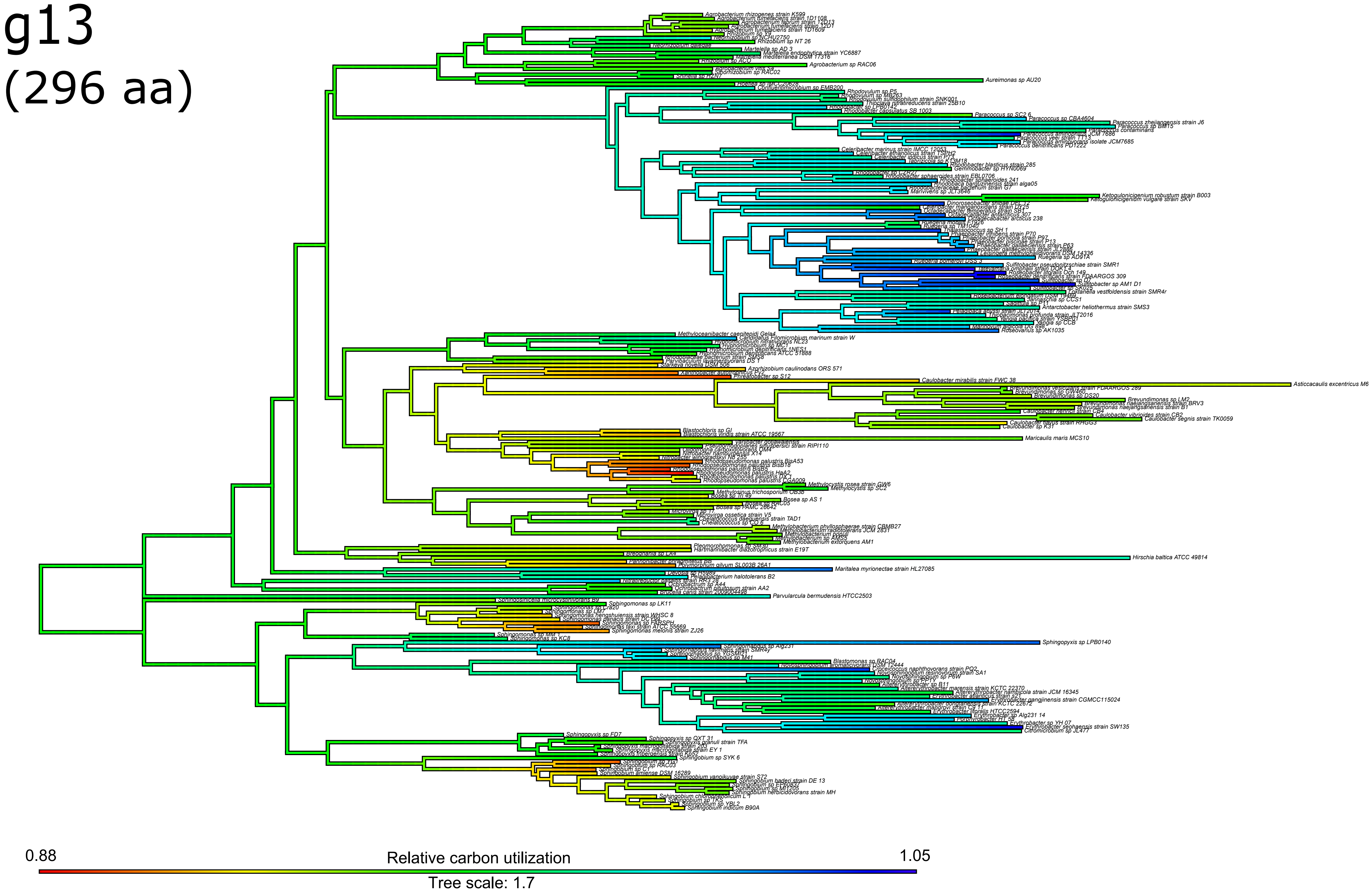
g13
(296 aa)
0.88
Relative carbon utilization
1.05
Tree scale: 1.7

# g14
# (150 aa)

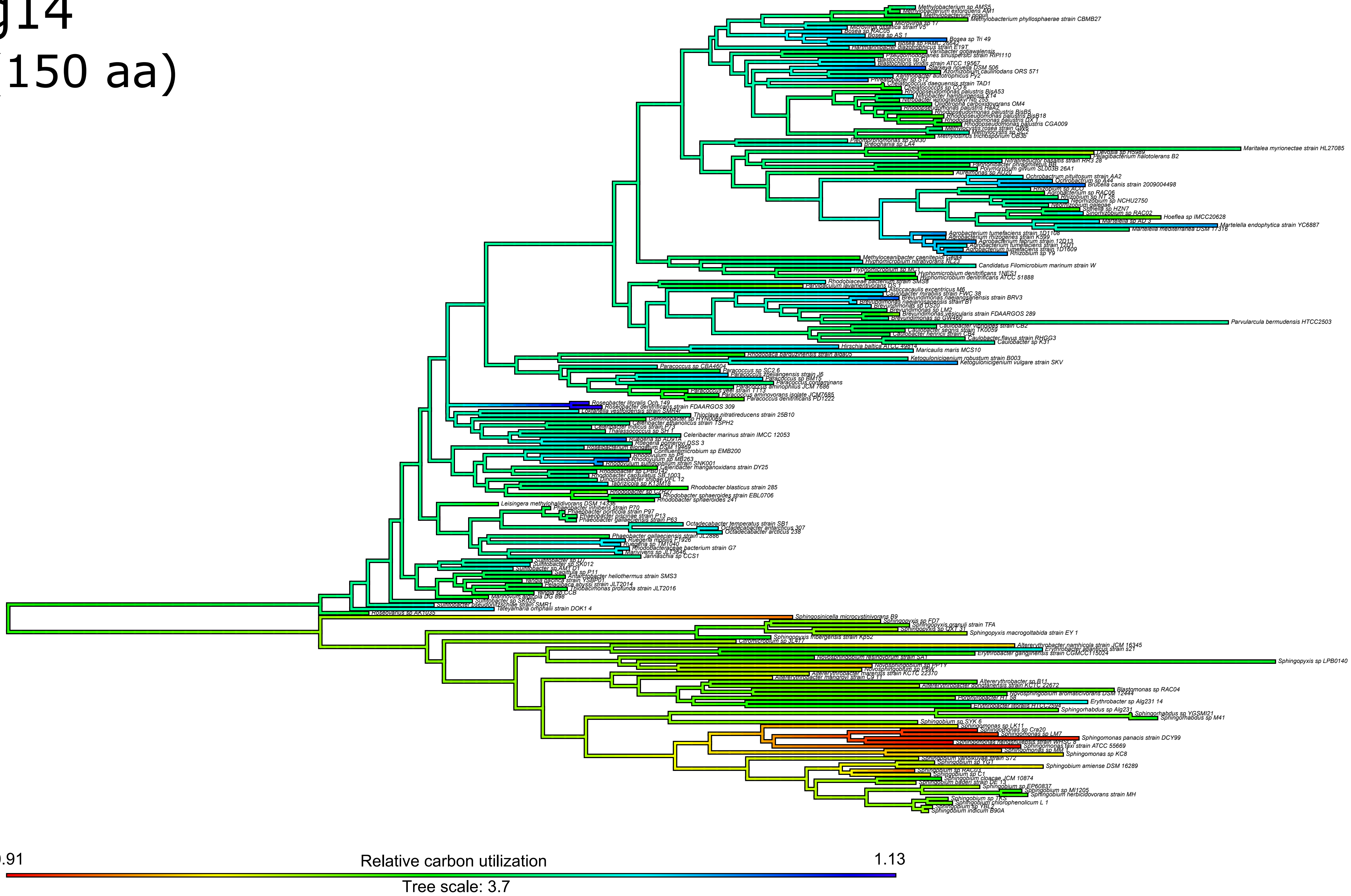

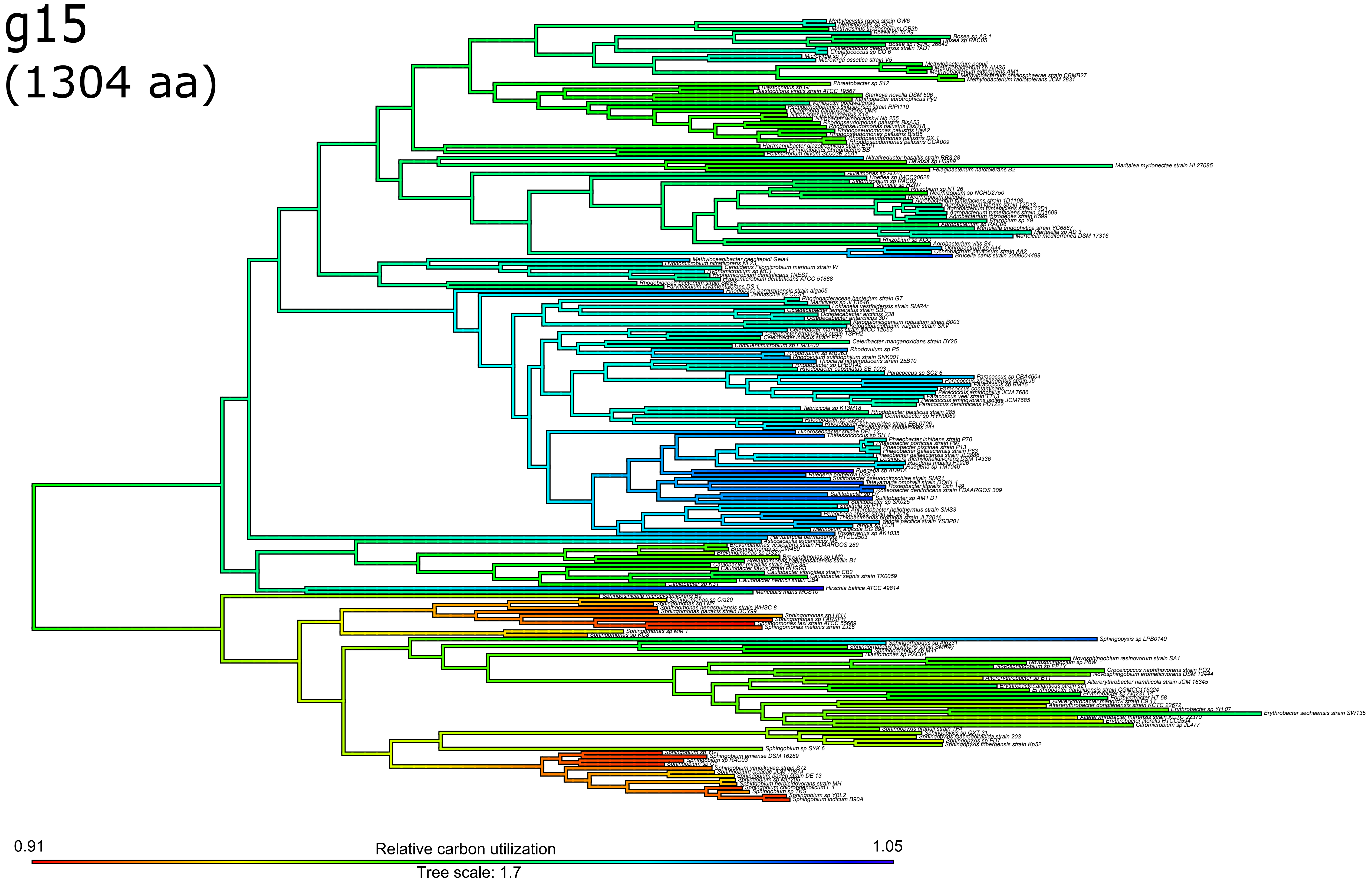
g15
(1304 aa)
0.91
Relative carbon utilization
1.05
Tree scale: 1.7
