## Supplemental Tables S1-S5 for "Selection for reducing energy cost of protein production drives the GC content and amino acid composition bias in gene transfer agents"

**Table S1. The number of detected GTA genes in 212 bacterial genomes and viruses from the RefSeq database.**

| GTA gene | Number of GTA genes | Number of viral homologs | Used for cost of production analysis | Used for ancestral reconstructions | Functional annotation[#] |
|---|---|---|---|---|---|
| *g1* | 41 | 0 | | | hypothetical protein |
| *g2* | 205 | 16 | X | X | ATP-binding protein |
| *g3* | 204 | 261 | X | X | phage-portal protein |
| *g3.5* | 57 | 0 | | | hypothetical protein |
| *g4* | 205 | 164 | X | X | HK97 family phage prohead protease |
| *g5* | 205 | 137 | X | X | phage major capsid protein |
| *g6* | 209 | 36 | X | X | hypothetical protein |
| *g7* | 151 | 63 | X | X | head-tail adaptor protein |
| *g8* | 209 | 66 | X | X | DUF3168 domain-containing protein |
| *g9* | 209 | 18 | X | X | phage major tail protein, TP901-1 family |
| *g10* | 208 | 1 | | X | gene transfer agent family protein |
| *g10.1* | 203 | 0 | | | phage tail assembly chaperone |
| *g11* | 209 | 1 | | X | phage tail tape measure protein |
| *g12* | 209 | 26 | X | X | TIGR02217 family protein |
| *g13* | 207 | 70 | X | X | DUF2163 domain-containing protein |
| *g14* | 200 | 26 | X | X | peptidase |
| *g15* | 200 | 12 | X | X | hypothetical protein |

[#] Taken from the RefSeq record NC_014034.1

**Table S2. Functional annotations of 26 single-copy genes that were used to calculate the normalized carbon utilization. Of these 26 genes,** 20 that belong to the "J" COG category were designated as "highly-expressed" genes.

| Gene ID | COG category | Putative Function# | UniProt Annotation |
|---|---|---|---|
| RPSH | J | One of the primary rRNA binding proteins, it binds directly to 16S rRNA central domain where it helps coordinate assembly of the platform of the 30S subunit. | 30S ribosomal protein S8 |
| RPSI | J | 30S ribosomal protein S9. | 30S ribosomal protein S9 |
| YCHF | J | GTP-dependent nucleic acid-binding protein engD. | Ribosome-binding ATPase YchF |
| TSF | J | Associates with the EF-Tu.GDP complex and induces the exchange of GDP to GTP. It remains bound to the aminoacyl-tRNA.EF- Tu.GTP complex up to the GTP hydrolysis stage on the ribosome. | Elongation factor Ts |
| INFC | J | IF-3 binds to the 30S ribosomal subunit and shifts the equilibrum between 70S ribosomes and their 50S and 30S subunits in favor of the free subunits, thus enhancing the availability of 30S subunits on which protein synthesis initiation begins. | Translation initiation factor IF-3 |
| FRR | J | Responsible for the release of ribosomes from messenger RNA at the termination of protein biosynthesis. May increase the efficiency of translation by recycling ribosomes from one round of translation to another. | Ribosome-recycling factor |
| RPSA | J | It is needed to translate mRNA with a short Shine-Dalgarno (SD) purine-rich sequence. | 30S ribosomal protein S1 |
| RSMA | J | Specifically dimethylates two adjacent adenosines (A1518 and A1519) in the loop of a conserved hairpin near the 3'-end of 16S rRNA in the 30S particle. May play a critical role in biogenesis of 30S subunits. | Ribosomal RNA small subunit methyltransferase A |
| RPSC | J | Binds the lower part of the 30S subunit head. Binds mRNA in the 70S ribosome, positioning it for translation. | 30S ribosomal protein S3 |
| RPLT | J | Binds directly to 23S ribosomal RNA and is necessary for the in vitro assembly process of the 50S ribosomal subunit. It is not involved in the protein synthesizing functions of that subunit. | 50S ribosomal protein L20 |

| | | | |
|---|---|---|---|
| RPLV | J | The globular domain of the protein is located near the polypeptide exit tunnel on the outside of the subunit, while an extended beta-hairpin is found that lines the wall of the exit tunnel in the center of the 70S ribosome. | 50S ribosomal protein L22 |
| RPLM | J | This protein is one of the early assembly proteins of the 50S ribosomal subunit, although it is not seen to bind rRNA by itself. It is important during the early stages of 50S assembly. | 50S ribosomal protein L13 |
| RPLO | J | Binds to the 23S rRNA. | 50S ribosomal protein L15 |
| RPLX | J | One of the proteins that surrounds the polypeptide exit tunnel on the outside of the subunit. | 50S ribosomal protein L24 |
| RPLB | J | One of the primary rRNA binding proteins. Required for association of the 30S and 50S subunits to form the 70S ribosome, for tRNA binding and peptide bond formation. It has been suggested to have peptidyltransferase activity. | 50S ribosomal protein L2 |
| RIMM | J | An accessory protein needed during the final step in the assembly of 30S ribosomal subunit, possibly for assembly of the head region. Probably interacts with S19. Essential for efficient processing of 16S rRNA. May be needed both before and after RbfA during the maturation of 16S rRNA. It has affinity for free ribosomal 30S subunits but not for 70S ribosomes. | Ribosome maturation factor RimM |
| RPLC | J | One of the primary rRNA binding proteins, it binds directly near the 3'-end of the 23S rRNA, where it nucleates assembly of the 50S subunit. | 50S ribosomal protein L3 |
| RPSK | J | Located on the platform of the 30S subunit, it bridges several disparate RNA helices of the 16S rRNA. Forms part of the Shine-Dalgarno cleft in the 70S ribosome. | 30S ribosomal protein S11 |
| RPLF | J | This protein binds to the 23S rRNA, and is important in its secondary structure. It is located near the subunit interface in the base of the L7 L12 stalk, and near the tRNA binding site of the peptidyltransferase center. | 50S ribosomal protein L6 |
| RPLD | J | One of the primary rRNA binding proteins, this protein initially binds near the 5'-end of the 23S rRNA. It is important during the | 50S ribosomal protein L4 |

| | | | |
|---|---|---|---|
| | | early stages of 50S assembly. It makes multiple contacts with different domains of the 23S rRNA in the assembled 50S subunit and ribosome. | |
| RNC | K | Digests double-stranded RNA. Involved in the processing of primary rRNA transcript to yield the immediate precursors to the large and small rRNAs (23S and 16S). Also processes some mRNAs, and tRNAs when they are encoded in the rRNA operon. | Ribonuclease 3 |
| RSMD | L | Methyltransferase | Ribosomal RNA small subunit methyltransferase D |
| RUVX | L | Could be a nuclease that resolves Holliday junction intermediates in genetic recombination. | Putative pre-16S rRNA nuclease |
| RECR | L | May play a role in DNA repair. It seems to be involved in an RecBC-independent recombinational process of DNA repair. It may act with RecF and RecO. | Recombination protein RecR |
| SMPB | O | Binds specifically to the SsrA RNA (tmRNA) and is required for stable association of SsrA with ribosomes. | SsrA-binding protein |
| GCP | O | Required for the formation of a threonylcarbamoyl group on adenosine at position 37 (t(6)A37) in tRNAs that read codons beginning with adenine. | Putative tRNA threonylcarbamoyladenosine biosynthesis protein Gcp |

[#]Based on the eggNOG-mapper annotation

**Table S3. The likelihood ratio test for the branch site A model.**

| GTA gene | LRT | p-value | Corrected p-value |
|---|---|---|---|
| ***g2*[$]** | **52.23** | **4.94E-13** | **6.91E-12** |
| ***g3*** | **58.29** | **2.26E-14** | **3.17E-13** |
| ***g4*** | **28.52** | **9.27E-08** | **1.30E-06** |
| ***g5*** | **33.89** | **5.83E-09** | **8.16E-08** |
| ***g6*** | **19.49** | **1.01E-05** | **0.00014112** |
| *g7* | 11.34 | 0.000759 | 0.0106204 |
| *g8* | 1.14 | 0.2857 | 1 |
| ***g9*** | **14.98** | **0.000109** | **0.0015218** |
| *g10* | 9.16 | 0.002474 | 0.034636 |
| *g11* | 6.09 | 0.01359 | 0.19026 |
| ***g12**** | **49.76** | **1.74E-12** | **2.43E-11** |
| ***g13*** | **51.84** | **6.02E-13** | **8.43E-12** |
| *g14* | 5.4 | 0.02014 | 0.28196 |
| ***g15**** | **153.36** | **3.20E-35** | **4.47E-34** |

*To reduce computation time, for *g12* and *g15* genes the alignments were trimmed by removing sites that had > 50% and >20% gaps, respectively.

[$]Genes in bold font have corrected p-values < 0.01

**Table S4. The average number of carbons per amino acid side chain in *Sphigomonadales* and three other orders combined together.**

| GTA gene | *Sphingomonadales* average | Others average | p-value | Corrected p-value |
|---|---|---|---|---|
| ***g2***[$] | **2.7** | **2.79** | **6.70E-17** | **9.38E-16** |
| ***g3*** | **2.71** | **2.81** | **3.80E-16** | **5.32E-15** |
| ***g4*** | **2.73** | **2.8** | **0.00017** | **0.00238** |
| ***g5*** | **2.65** | **2.75** | **1.80E-21** | **2.52E-20** |
| ***g6*** | **2.72** | **2.82** | **2.20E-09** | **3.08E-08** |
| *g7* | 2.79 | 2.79 | 0.47 | 1 |
| *g8* | 2.65 | 2.71 | 0.99 | 1 |
| ***g9*** | **2.53** | **2.67** | **4.40E-22** | **6.16E-21** |
| *g10* | 2.67 | 2.46 | 1 | 1 |
| *g11* | 2.34 | 2.37 | 0.004 | 0.056 |
| ***g12*** | **2.74** | **2.86** | **5.10E-15** | **7.14E-14** |
| *g13* | 2.63 | 2.68 | 0.003 | 0.042 |
| *g14* | 2.69 | 2.89 | 7.80E-19 | 1.09E-17 |
| ***g15*** | **2.62** | **2.71** | **2.96E-11** | **4.15E-10** |

[$]Genes in bold font have sites under the positive selections and corrected p-values < 0.01

**Table S5. Contribution of sites under the positive selection to the carbon utilization in *Sphingomonadales'* GTA genes.**

| Site | Bayes empirical Bayes (probability) | Average change in number of carbons | p-value |
|---|---|---|---|
| g2 | | | |
| 361 | 1 | +6.00 | 1.0000 |
| 433 | 1 | 0.00 | 1.0000 |
| 466 | 0.95 | +0.26 | 0.9856 |
| **479$** | **1** | **-2.20** | **0.0000** |
| **503** | **1** | **-0.93** | **0.0000** |
| 505 | 1 | +0.10 | 0.9940 |
| **506** | **1** | **-3.39** | **0.0000** |
| **683** | **0.98** | **-1.16** | **0.0000** |
| **759** | **0.99** | **-1.31** | **0.0000** |
| 762 | 0.99 | +0.18 | 0.8826 |
| **812** | **0.98** | **-2.33** | **0.0000** |
| 815 | 1 | +1.90 | 1.0000 |
| 821 | 0.97 | 0.00 | 1.0000 |
| g3 | | | |
| **127** | **0.962** | **-0.74** | **0.0000** |
| 140 | 0.974 | +3.26 | 1.0000 |
| **142** | **0.95** | **-1.29** | **0.0000** |
| 188 | 0.985 | +0.36 | 0.9127 |
| **193** | **0.988** | **-1.55** | **0.0000** |
| 199 | 0.964 | +0.92 | 1.0000 |
| **205** | **0.995** | **-2.98** | **0.0000** |
| **211** | **0.969** | **-0.72** | **0.0000** |
| **216** | **0.952** | **-0.66** | **0.0000** |
| **245** | **0.971** | **-0.75** | **0.0000** |
| **247** | **0.995** | **-1.70** | **0.0000** |
| **248** | **0.972** | **-0.89** | **0.0000** |
| **249** | **0.984** | **-0.88** | **0.0000** |
| **256** | **0.992** | **-1.34** | **0.0000** |
| **262** | **0.999** | **-5.53** | **0.0000** |
| 268 | 0.993 | +2.82 | 1.0000 |
| 274 | 0.967 | +0.06 | 0.1559 |
| 349 | 0.973 | +2.75 | 1.0000 |
| **375** | **0.971** | **-2.22** | **0.0000** |
| **376** | **0.961** | **-1.08** | **0.0000** |
| **386** | **0.955** | **-0.96** | **0.0000** |

| | | | |
|---|---|---|---|
| **396** | **0.97** | **-0.36** | **0.0000** |
| 400 | 0.952 | +0.19 | 1.0000 |
| **438** | **0.953** | **-1.23** | **0.0000** |
| 454 | 0.993 | +1.03 | 1.0000 |
| **456** | **0.98** | **-0.68** | **0.0000** |
| **469** | **0.968** | **-1.84** | **0.0000** |
| 491 | 0.989 | +0.02 | 0.1556 |
| **507** | **0.975** | **-2.65** | **0.0000** |
| 520 | 0.954 | +2.31 | 1.0000 |
| **526** | **0.973** | **-8.57** | **0.0000** |
| **572** | **0.992** | **-0.34** | **0.0183** |
| 586 | 0.979 | +1.57 | 1.0000 |
| g4 | | | |
| 110 | 0.969 | +1.52 | 0.9999 |
| 115 | 1 | 0.00 | 1.0000 |
| **116** | **0.958** | **-0.26** | **0.0000** |
| 122 | 0.964 | -0.35 | 0.6569 |
| 128 | 0.95 | +1.34 | 1.0000 |
| 135 | 0.951 | +0.71 | 1.0000 |
| 151 | 0.971 | +0.26 | 0.9976 |
| **152** | **0.989** | **-0.88** | **0.0000** |
| 153 | 0.986 | +0.91 | 1.0000 |
| 165 | 0.951 | +1.07 | 0.7478 |
| **166** | **1** | **-5.50** | **0.0000** |
| **169** | **0.961** | **-0.87** | **0.0000** |
| **177** | **1** | **-2.76** | **0.0000** |
| 179 | 0.97 | +0.09 | 0.6501 |
| **183** | **0.99** | **-1.52** | **0.0000** |
| **184** | **0.951** | **-0.57** | **0.0006** |
| 185 | 0.952 | +0.70 | 1.0000 |
| **186** | **0.974** | **-0.72** | **0.0000** |
| **188** | **0.994** | **-1.82** | **0.0000** |
| 196 | 0.989 | +0.30 | 0.9984 |
| 206 | 0.993 | +2.71 | 1.0000 |
| 208 | 0.994 | 0.00 | 1.0000 |
| 210 | 0.999 | +0.48 | 1.0000 |
| 211 | 0.993 | +0.53 | 1.0000 |
| **227** | **0.976** | **-1.86** | **0.0000** |
| **238** | **0.984** | **-0.66** | **0.0000** |
| **245** | **1** | **-5.29** | **0.0000** |
| **255** | **0.951** | **-0.77** | **0.0000** |
| 257 | 0.999 | +1.16 | 1.0000 |
| g5 | | | |

| | | | |
|---|---|---|---|
| 413 | 0.981 | +5.22 | 1.0000 |
| **414** | **0.976** | **-3.44** | **0.0000** |
| **450** | **0.954** | **-2.88** | **0.0000** |
| **524** | **0.986** | **-1.08** | **0.0000** |
| **586** | **0.992** | **-1.40** | **0.0000** |
| **588** | **0.952** | **-1.04** | **0.0000** |
| 590 | 0.988 | +2.99 | 1.0000 |
| **614** | **0.982** | **-3.43** | **0.0000** |
| **623** | **0.98** | **-1.93** | **0.0000** |
| 635 | 0.973 | +1.15 | 1.0000 |
| 710 | 0.976 | +1.41 | 1.0000 |
| **712** | **0.977** | **-0.27** | **0.0000** |
| g6 | | | |
| **103** | **0.979** | **-2.51** | **0.0000** |
| **118** | **0.971** | **-2.21** | **0.0000** |
| 178 | 0.956 | -0.06 | 0.1520 |
| **188** | **0.991** | **-0.66** | **0.0000** |
| 192 | 0.98 | +0.41 | 1.0000 |
| 227 | 0.953 | +1.11 | 1.0000 |
| **231** | **0.986** | **-2.57** | **0.0000** |
| **281** | **0.991** | **-0.43** | **0.0000** |
| 289 | 0.999 | +5.36 | 1.0000 |
| 290 | 0.968 | +1.99 | 1.0000 |
| 292 | 0.969 | +1.35 | 1.0000 |
| g9 | | | |
| **7** | **1** | **-2.93** | **0.0000** |
| **8** | **0.992** | **-0.97** | **0.0000** |
| 27 | 0.993 | +0.03 | 0.6380 |
| **36** | **0.962** | **-1.11** | **0.0000** |
| **38** | **0.995** | **-0.39** | **0.0000** |
| **40** | **0.982** | **-3.00** | **0.0000** |
| **42** | **0.99** | **-1.01** | **0.0000** |
| 46 | 0.992 | +0.33 | 1.0000 |
| 50 | 1 | +1.81 | 1.0000 |
| **65** | **0.986** | **-0.24** | **0.0000** |
| **67** | **1** | **-1.19** | **0.0000** |
| 72 | 0.988 | +0.65 | 1.0000 |
| **77** | **0.983** | **-1.30** | **0.0000** |
| **78** | **0.988** | **-1.96** | **0.0000** |
| **79** | **0.997** | **-0.72** | **0.0000** |
| **84** | **0.982** | **-0.65** | **0.0002** |
| **97** | **0.991** | **+0.04** | **0.0012** |
| **100** | **0.985** | **-2.41** | **0.0000** |

| | | | |
|---|---|---|---|
| 102 | 1 | 0.00 | 1.0000 |
| **104** | **1** | **-6.96** | **0.0000** |
| 105 | 0.986 | +2.68 | 1.0000 |
| 108 | 0.995 | +0.88 | 1.0000 |
| 110 | 0.994 | +1.05 | 1.0000 |
| 114 | 1 | +0.51 | 1.0000 |
| 126 | 0.998 | +1.97 | 1.0000 |
| 131 | 0.998 | +0.53 | 1.0000 |
| 134 | 1 | 0.00 | 1.0000 |
| 135 | 0.985 | +0.03 | 0.9122 |
| **140** | **1** | **-5.41** | **0.0000** |
| g12 | | | |
| **6** | **0.955** | **-1.09** | **0.0000** |
| **13** | **0.988** | **-1.87** | **0.0000** |
| 21 | 0.999 | +2.54 | 1.0000 |
| **22** | **1** | **-2.79** | **0.0000** |
| **28** | **0.999** | **-2.71** | **0.0000** |
| **38** | **0.98** | **-0.77** | **0.0000** |
| **44** | **0.992** | **-0.49** | **0.0000** |
| **55** | **1** | **-1.02** | **0.0000** |
| **59** | **0.997** | **-1.61** | **0.0000** |
| 66 | 1 | +1.00 | 1.0000 |
| **72** | **0.999** | **-2.97** | **0.0000** |
| 86 | 0.967 | +0.81 | 1.0000 |
| 88 | 0.999 | +0.02 | 0.5324 |
| 89 | 0.965 | +0.55 | 0.9976 |
| 99 | 0.981 | +0.01 | 0.8037 |
| 110 | 0.97 | -0.39 | 0.0107 |
| **122** | **0.982** | **-3.80** | **0.0000** |
| 125 | 0.97 | +0.67 | 1.0000 |
| 149 | 1 | +1.76 | 1.0000 |
| **152** | **0.983** | **-0.32** | **0.0014** |
| 155 | 0.998 | +0.06 | 0.7210 |
| 184 | 0.999 | +0.82 | 1.0000 |
| **202** | **0.991** | **-0.37** | **0.0000** |
| g13 | | | |
| 18 | 0.97 | +2.22 | 1.0000 |
| 27 | 0.996 | +0.04 | 0.9421 |
| 33 | 0.991 | +0.69 | 0.9986 |
| **44** | **0.985** | **-0.94** | **0.0000** |
| **69** | **0.967** | **-1.95** | **0.0000** |
| 70 | 0.969 | +1.30 | 1.0000 |
| 71 | 0.984 | +1.96 | 1.0000 |

| | | | |
|---|---|---|---|
| **72** | **0.998** | **-0.86** | **0.0002** |
| 76 | 0.974 | +0.82 | 1.0000 |
| **78** | **0.967** | **-0.59** | **0.0028** |
| 86 | 0.991 | +0.60 | 0.9994 |
| 94 | 0.991 | +0.41 | 0.4857 |
| 114 | 0.98 | +1.09 | 1.0000 |
| **131** | **0.984** | **-0.22** | **0.0000** |
| **135** | **0.991** | **-1.80** | **0.0000** |
| **143** | **0.964** | **-0.98** | **0.0000** |
| **170** | **0.959** | **-1.00** | **0.0000** |
| **173** | **0.965** | **-1.45** | **0.0000** |
| **174** | **0.977** | **-0.35** | **0.0000** |
| **175** | **0.968** | **-0.66** | **0.0001** |
| 177 | 0.995 | +2.47 | 1.0000 |
| 186 | 0.97 | +1.14 | 0.9999 |
| **231** | **0.99** | **-5.97** | **0.0000** |
| 238 | 0.979 | +1.68 | 0.9999 |
| 242 | 0.988 | +1.42 | 1.0000 |
| 244 | 0.972 | +1.42 | 1.0000 |
| **284** | **0.966** | **-6.68** | **0.0000** |
| **301** | **0.964** | **-2.67** | **0.0000** |
| 302 | 0.977 | +0.47 | 0.9735 |
| **327** | **0.999** | **-3.96** | **0.0000** |
| **337** | **0.967** | **-1.78** | **0.0000** |
| g15 | | | |
| 37 | 0.957 | +2.34 | 1.0000 |
| 54 | 0.998 | +4.06 | 1.0000 |
| **76** | **0.954** | **-1.25** | **0.0000** |
| 107 | 0.987 | +0.04 | 0.9073 |
| 125 | 0.981 | +2.26 | 1.0000 |
| **126** | **0.985** | **-3.30** | **0.0000** |
| 170 | 0.961 | -0.10 | 0.1523 |
| **187** | **0.966** | **-2.96** | **0.0000** |
| 205 | 0.956 | +1.56 | 1.0000 |
| **219** | **0.974** | **-1.75** | **0.0000** |
| **222** | **0.95** | **-1.23** | **0.0000** |
| 302 | 0.984 | +0.87 | 1.0000 |
| 376 | 0.983 | +0.24 | 0.8946 |
| **403** | **0.962** | **-1.96** | **0.0000** |
| **408** | **0.955** | **-1.06** | **0.0000** |
| 413 | 0.961 | +0.91 | 0.9775 |
| **422** | **0.965** | **-0.78** | **0.0000** |
| **427** | **0.957** | **-0.63** | **0.0020** |

| | | | |
|---|---|---|---|
| **435** | **0.987** | **-2.52** | **0.0000** |
| 509 | 0.953 | +2.58 | 1.0000 |
| **512** | **0.975** | **-6.30** | **0.0000** |
| 547 | 0.954 | -1.77 | 0.0120 |
| 551 | 0.957 | +1.18 | 1.0000 |
| **574** | **0.968** | **-4.20** | **0.0000** |
| **622** | **0.957** | **-0.52** | **0.0000** |
| **624** | **0.952** | **-2.42** | **0.0000** |
| 684 | 0.987 | +1.93 | 1.0000 |

[$]Sites in bold font significantly reduce carbon utilization in *Sphingomonadales* with corrected p-values < 0.01
